# An improved FLARE system for recording and manipulating neuronal activity

**DOI:** 10.1101/2025.01.13.632875

**Authors:** Guanwei Zhou, Ruonan Li, Ola Bartolik, Yuqian Ma, Wei Wei Wan, Jennifer Meng, Yujia Hu, Bing Ye, Wenjing Wang

**Author notes:** These authors contributed equally (Guanwei Zhou, Ruonan Li). Correspondence (W.W.), (B. Y.).

## Abstract

Recording and manipulating neuronal ensembles that underlie cognition and behavior *in vivo* is challenging. FLARE is a light– and calcium-gated transcriptional reporting system for labeling activated neurons on the order of minutes. However, FLARE is limited by its sensitivity to prolonged neuronal activities. Here, we present an improved version of FLARE, termed cytoFLARE. cytoFLARE incorporates cytosolic expression of the transcription factor and a more sensitive pair of calcium sensing domains. We showed that cytoFLARE provides more calcium– and light-dependent signals in HEK293T cells and higher signal-to-background ratios in neuronal cultures. We further established cytoFLARE transgenic *Drosophila* models and applied cytoFLARE to label activated neurons upon sensory or optogenetic stimulation within a defined time window. Notably, through cytoFLARE-driven expression of an optogenetic actuator, we successfully reactivated neurons involved in the larval nociceptive system. Our findings demonstrate the first characterization and application of time-gated calcium integrators in *Drosophila*.

## Introduction

A central challenge in neuroscience is the ability to record and manipulate neuronal ensembles that underlie specific cognitive and behavioral processes. During a specific time window of these processes, only a subset of neuronal ensembles are activated. For example, a subset of hippocampal CA1 cells is activated when an animal experiences a novel context. The same subset of neurons will be re-activated when the animal returns to the same context but not to a different novel context ^1^. Labeling and manipulating such ensembles are essential for studying mechanisms underlying cognition and behavior ^2^. To identify these activated neurons from the large population of neurons sharing similar morphology and genetic profiles, a variety of genetically encoded tools for labeling and manipulating neurons based on the neuronal activity have been developed. Among the tools for recording neuronal activity, real-time imaging has played a crucial role in monitoring dynamic calcium levels ^3-5^ and voltage changes ^6-8^, with subsecond precision. However, real-time imaging is restricted to small fields of view in deep tissues and does not allow further characterizations of the neuronal population based on the activity detected, including manipulating their activities for investigating their roles in behaviors.

Neuronal activity-dependent expression of immediate early genes (IEGs) such as c-Fos ^1^, Arc ^9^, or their derivatives ^10^, offer an approach for addressing the limitation of real-time imaging. Time-gated IEG systems, such as TRAP ^11^, TetTag ^12^, and CANE^13^, were further developed to label active neuronal subpopulations within a specific time window. Through transgenic expression of fluorescent proteins, IEG-based approaches provide large-scale “snapshots” of activated neurons. IEG enhancers can also be used to express optogenetic or chemogenetic actuators for interrogating the causal relationship between the targeted neural circuit and a cognitive or behavioral process ^14-16^. However, since activity-dependent expression of IEGs varies widely across neuron types and brain areas ^17,18^, findings from IEG-based reporters are often difficult to interpret and generalize to different circuits. Moreover, the poor temporal resolution, typically ranging from hours to days, limits IEG-based tools for studying temporally precise cognitive and behavioral processes. Additionally, IEG-based reporters are not currently applicable for all experimental organisms, such as *Drosophila*, due to the lack of appropriate IEGs. Instead, calcium-dependent transcriptional reporters, such as CaLexA ^19^ and TRIC ^20^, were developed for studying neuronal activity in *Drosophila*. However, they also report slow changes in neuronal activity without time-gating.

Temporally-gated calcium integrators have been developed to improve the temporal resolution. For example, CaMPARI is a calcium-dependent photoconvertible fluorescent protein and can permanently label activated neurons with subsecond resolution ^21,22^. However, CaMPARI requires short-wavelength light (405nm), limiting its utility. To overcome this drawback, a blue light– and calcium-gated enzymatic reporter, LiPPI-CLAPon, was developed, but it has not been applied in neuronal culture or *in vivo* yet ^23^. Neither CaMPARI nor LiPPI-CLAPon allow activity-based neural circuit manipulation.

Blue light– and calcium-gated transcriptional reporters have been developed to provide more versatile readout with high temporal precision ^24-29^. In FLARE ^24^ and its more advanced variant FLiCRE ^28^, a transmembrane-bound transcription factor can translocate to the nucleus only upon protease cleavage in the coincidence of both light and high calcium. In these systems, blue light is used to activate the light, oxygen, and voltage-sensing (LOV) domain to uncage the protease cleavage site that links a transcription factor to the transmembrane domain. At an increased level of intracellular calcium that coincides with blue light, the calmodulin (CaM)-linked protease (TEVp) is recruited to the proximity of the protease cleavage site (TEVcs) via the calcium-dependent interaction between CaM and the CaM-binding peptide (MK2), leading to the cleavage of the TEVcs and thus the release of the transcription factor. The transcription factor can then enter the nucleus to induce the transcription of reporters or optogenetic or chemogenetic actuators. Compared to a conceptually similar reporter Cal-Light ^25,29^, which utilizes split TEVp, FLARE series reporters have faster kinetics. In contrast to existing IEG-based reporters, FLARE provides improved temporal resolution on the order of seconds to minutes. However, current FLARE reporters are still limited by their low sensitivity to transient calcium spikes. In addition, they have not yet been demonstrated in *Drosophila*, which lacks time-gated transcriptional systems for reporting and manipulating neuronal activity during cognitive and behavioral processes.

Here, we report cytoFLARE (cytosol-expressed FLARE), an improved variant of FLARE. Through cytosolic expression of transcription factor and a more sensitive pair of CaM-MK2, cytoFLARE shows 2.7-fold improved calcium– and light-dependent signal, compared to the published FLiCRE for detecting transient calcium elevation in HEK293T cells and 1.8-fold improved signal-to-background ratio (SBR) in primary rat neurons. Furthermore, we establish cytoFLARE transgenic *Drosophila* models for detecting calcium increase on the order of minutes. We also demonstrate that cytoFLARE can not only record transient calcium increase in response to physiological and optogenetic stimuli, but also manipulate neural ensembles involved in behavior of *Drosophila* larvae.

## Results

### cytoFLARE engineering and characterization in heterologous cell cultures

In the cytoFLARE design, we first replaced the transmembrane domain, which excludes the transcription factor (TF) construct from the nucleus, with a cytosolic domain that prevents the TF construct entering the nucleus. The rationale behind this design is that a cytosol-located TF can more readily form a calcium-dependent complex with the TEV protease in the cytosol, facilitating the protease-cleavage-dependent release of the TF into the nucleus. Similarly, a recently developed soma-targeted Cal-Light showed enhanced labeling efficiency by increasing the chance of the two components binding ^29^. To localize the TF in the cytosol, we fused the TF construct with a supraphysiological nuclear export signal peptide (SNES3) ^30,31^ or a mutant estrogen ligand-binding domain (ER^T2^) ^32^ (Supplementary Fig. 1). ER^T2^ efficiently excluded the TF from the nucleus, showing low background transcriptional activation in the basal state and blue light– and high calcium-dependent activation of reporter gene expression. Therefore, we used ER^T2^ in the cytoFLARE design (Figure 1A and 1B).

**Figure 1.**
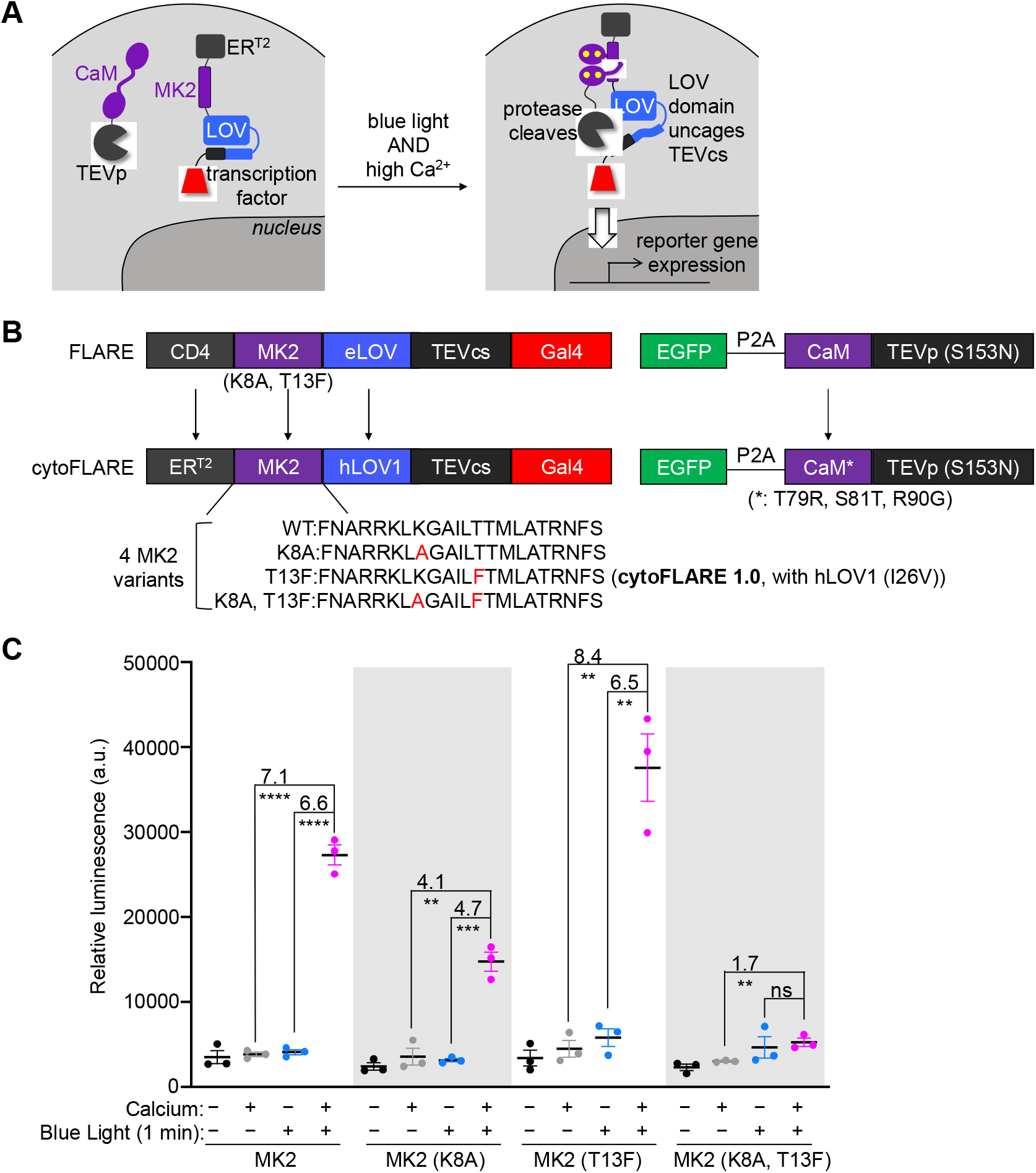
Design and optimization of cytoFLARE1.0. (A) Schematics of cytoFLARE design. ER^T2^ excludes the transcription factor from entering the nucleus in the absence of blue light and high calcium concentration. In the high calcium state, CaM and MK2 interact, bringing the TEVp into proximity with TEVcs, which can be unblocked from a LOV domain with blue light illumination. (B) Construct design of cytoFLARE, in comparison to FLARE. The arrows indicate the difference between the cytoFLARE and FLARE deisgn. (C) Optimization of cytoFLARE by screening four MK2 variants shown in panel B. HEK293T cells expressing cytoFLARE constructs were stimulated by blue light and high calcium for 1 minute. Luciferase was used as the reporter gene. Values above the dots represent SBRs which were calculated by dividing the means of the conditions indicated. Error bars, standard error of the mean. The mean is represented by the thicker horizontal bar. Stars represent significance after an unpaired two-tailed Student’s t-test. n = 3. ****p value < 0.0001; ***p value < 0.001; **p value < 0.01; ns, p value > 0.05.

To improve cytoFLARE’s sensitivity to changes in calcium levels, we introduced mutations from GCaMP6s ^4^ to the CaM in the cytoFLARE for improved calcium affinity. We then screened four variants of MK2 with different affinities for the calcium-bound CaM ^20,24^ with 1-minute light and calcium stimulation (Figure 1B). K8A mutation was previously incorporated to the MK2 in FLARE to reduce the binding of MK2 with CaM at the basal calcium level ^20^, and T13F was introduced to MK2 to achieve orthogonality with endogenous pair of CaM using a bump & hole strategy ^24^. We tested MK2 with or without K8A or T13F mutation in cytoFLARE constructs. Removal of K8A in the MK2 significantly enhanced the light– and calcium-dependent signal and maintained low background signals in the basal state (Figure 1C). To minimize interference from endogenous CaM, we chose to use the MK2 (T13F) in the cytoFLARE design, with light-dependent SBR (the ratio of the mean activation signal to the mean background signal) of 8.4 and calcium-dependent SBR of 6.5.

To further improve cytoFLARE’s light-dependent SBRs and activation efficiency, we next tested different LOV domains and TEVp. We previously replaced eLOV domain in FLARE by hLOV1 domain ^33^, which blocks TEVcs more efficiently in the dark, in the above screening (Figure 1B). We subsequently screened hLOV1 domains with different thermal reversion kinetics ^34^. As shown in Supplementary Fig. 2, hLOV1 provides the highest SBRs in cytoFLARE, followed by hLOV1 (I26V), and hLOV1 (V15T) upon 1-minute light and calcium stimulation. However, hLOV1 (I26V) enables faster reset of the light-stimulated LOV domain in the dark ^34^, providing faster temporal resolution. We thus chose to include hLOV1 (I26V) in the following cytoFLARE designs. We also screened TEVp with different kinetics ^26^. TEVp (T30A, S153N) resulted in higher background, while TEVp (T30A) and TEVp (S153N) provided similar SBRs in cytoFLARE (Supplementary Fig. 3). Therefore, we proceeded using TEVp (S153N) in cytoFLARE and named the cytoFLARE variant with MK2 (T13F), hLOV1 (I26V) and TEVp (S153N) as cytoFLARE1.0.

To evaluate the performance of cytoFLARE1.0, we then compared cytoFLARE1.0 with FLiCRE in HEK293T cells. For a more extensive evaluation, we also made FLiCRE2 by incorporating the optimal components from cytoFLARE1.0 (Supplementary Fig. 4). Both cytoFLARE1.0 and FLiCRE2 provided significantly higher signals than FLiCRE, upon 1-minute light and high calcium stimulation. However, FLiCRE2 led to higher background than cytoFLARE1.0 in the calcium only condition. Therefore, we moved forward with cytoFLARE1.0 for further characterizations.

Motivated by the improved calcium– and light-dependent signal of cytoFLARE1.0, we further compared cytoFLARE1.0 with FLiCRE in HEK293T cells. To investigate the temporal sensitivity of cytoFLARE, we characterized cytoFLARE1.0 with both 1-minute and 5-minute light and calcium stimulation using either the GAL4-driven mCherry (Figure 2A and 2B, Supplementary Fig. 5) or a luciferase reporter (Supplementary Fig. 6). cytoFLARE1.0 provided higher light– and calcium-dependent reporter activation with 5-minute stimulation than 1-minute stimulation. With both mCherry and luciferase reporter genes, cytoFLARE1.0 showed ∼2-fold higher light– and calcium-dependent reporter expression signal than FLiCRE under both 1-minute and 5-minute stimulation conditions. However, cytoFLARE1.0 did not always result in a significant increase in the SBRs, particularly with 5-minute stimulation due to increased background.

**Figure 2.**
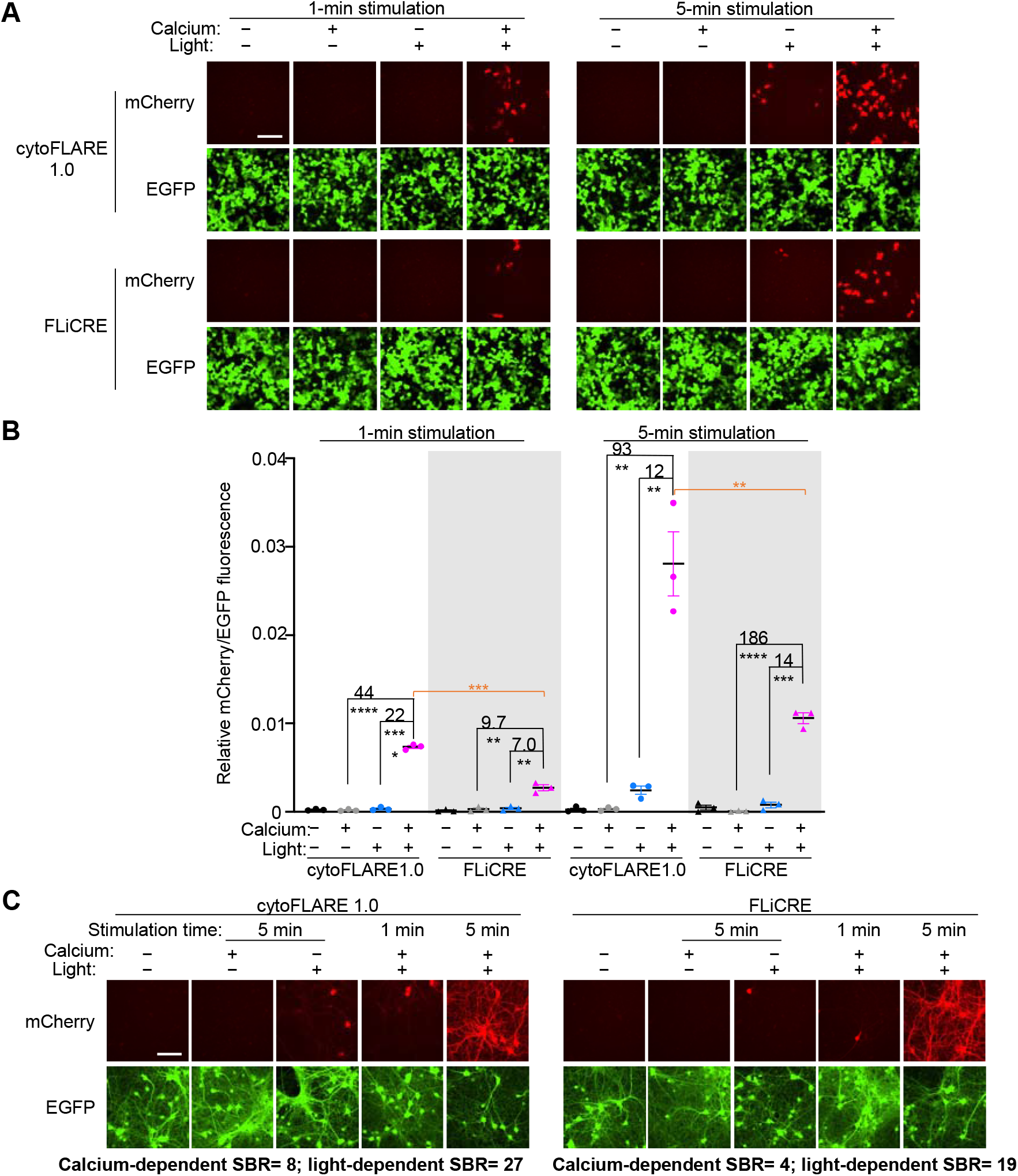
Characterization of cytoFLARE1.0 in HEK293T cells and primary rat neurons, in parallel with FLiCRE. (A) Representative fluorescence images for characterizing cytoFLARE1.0 in HEK293T cells in response to 1-minute and 5-minute blue light and high calcium stimulation. FLiCRE was tested in parallel for comparisons. mCherry, reporter gene expression. EGFP, expression marker of the TEVp construct. Scale bar, 100 μm. (B) Quantification of experiments in panel A. Values above the dots represent SBRs which were calculated by dividing the means of the conditions indicated. Error bars, standard error of the mean. The mean is represented by the thicker horizontal bar. Stars represent significance after an unpaired two-tailed Student’s t-test. n = 3. ****p value <0.0001; ***p value <0.001; **p value <0.01. (C) Representative fluorescence images for characterizing cytoFLARE1.0 and FLiCRE in primary rat neurons. Neurons were treated by bicuculline continuously for 1 and 5 minutes. Simultaneously, blue light was applied in pulses at 2 s on/ 4 s off for a total time of 1, 3, and 5 minutes. mCherry, reporter gene expression. EGFP, expression marker of the TEVp construct. Scale bar, 100 μm.

We then performed temporal sensitivity characterizations and comparisons of FLARE-based reporters with the tTA-driven mCherry reporter gene in primary rat neuronal culture. To induce calcium increase in the neurons, we treated cells with the GABA receptor antagonist, bicuculline, for 1 and 5 minutes, and simultaneously applied pulsed blue light (2s on/ 4s off). cytoFLARE1.0 provided higher SBRs than FLiCRE upon 5-minute stimulation (Figure 2C, Supplementary Fig. 7). Both cytoFLARE1.0 and FLiCRE failed to be significantly activated upon 1-minute stimulation. Therefore, the FLARE system is less sensitive in neuronal cultures than in HEK293T cells. To potentially further improve cytoFLARE sensitivity, we tested cytoFLARE1.1, which consists of the same TF construct as cytoFLARE1.0 but uses CaM-TEVp (T30A, S153N) to enable faster protease kinetics. cytoFLARE1.1 showed marginal improvement in activation compared to cytoFLARE1.0 (Supplementary Fig. 7).

### Establish cytoFLARE for recording neuronal activity during behavior in *Drosophila*

We generated transgenic *Drosophila* lines that express the two components of cytoFLARE1.0, ER^T2^-tethered TF (LexA) and CaM-TEVp (Figure 3A). Both components were driven by the UAS promoter, and these UAS-transgenic lines can be crossed with GAL4 driver lines to express cytoFLARE in select types of neurons. To report the LexA transcription activity, a LexAop2-mCherry transgene was introduced into the final flies (Figure 3A).

**Figure 3.**
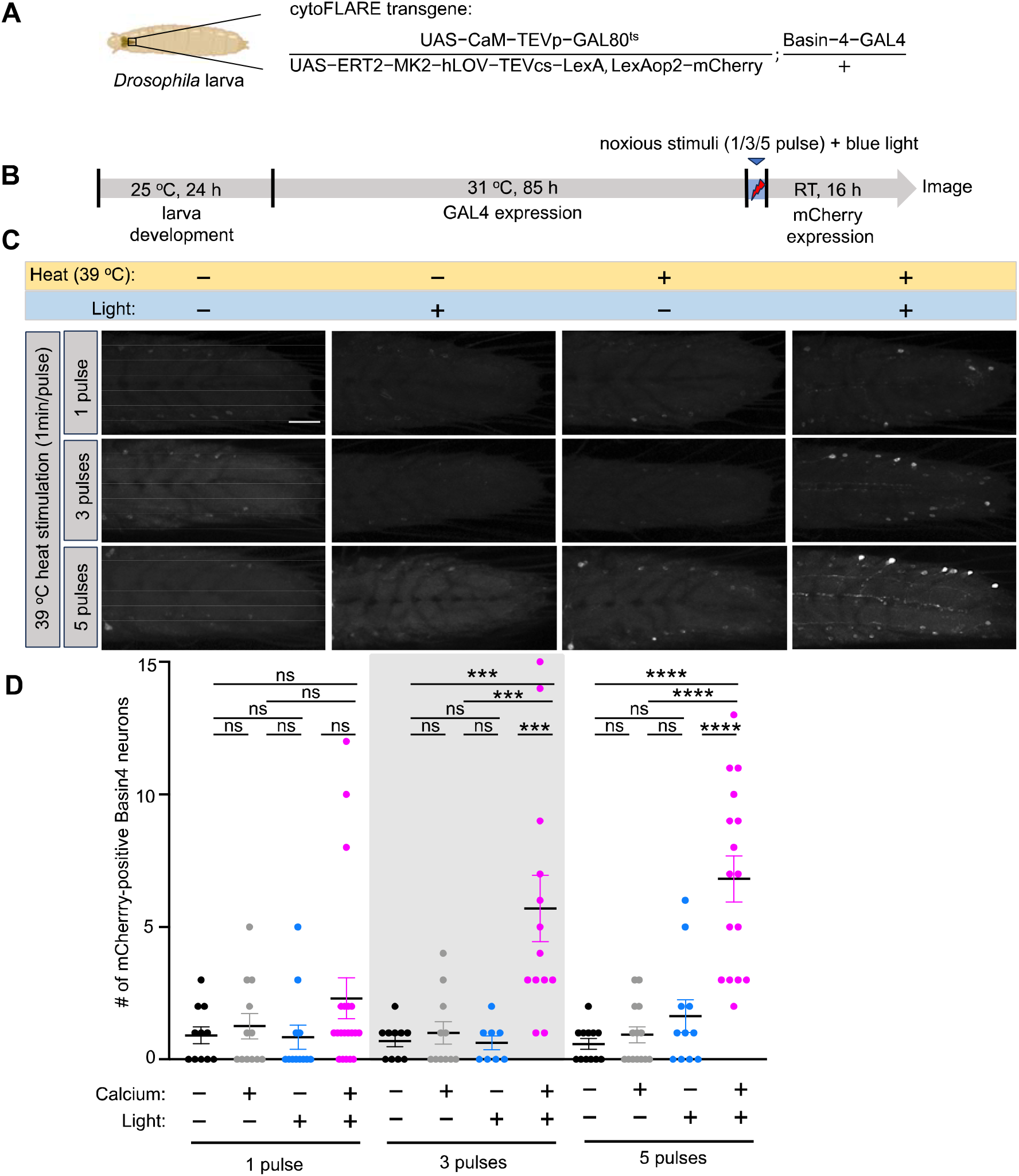
In vivo characterization of cytoFLARE1.0 in response to noxious stimuli in *Drosophila*. (A) Schematic of a larva expressing cytoFLARE1.0 in Basin-4 neurons with a temperature-sensitive Gal80 transgene. (B) Experimental workflow for characterizing cytoFLARE1.0 in response to noxious stimuli in Drosophila. After egg collection for 24 hours, larvae were kept at 31°C for 85 h to inactivate GAL80^ts^ so that the TEVp and ER^T2^-transcription factor transgenes could be expressed. The larvae were then exposed to 1, 3, or 5 pulses of noxious heat stimulation (39°C, 1 min on, 5 min off) and blue light (450 nm, 100 μW/mm^2^) or one of the two treatments. Subsequently, the larvae were kept at room temperature for 16 h to allow the expression of the mCherry fluorescent reporter. (C) Representative fluorescence images of Basin-4 neurons expressing mCherry, which indicates the cytoFLARE1.0 activation, in the condition shown in panel B. We have 4 groups: (1) with neither noxious stimulation nor TEVcs uncaging (-heat,-light); (2) with noxious stimulation but not TEVcs uncaging (+heat, −light); (3) without noxious stimulation but with TEVcs uncaging (-heat,+light); and (4) with both noxious stimulation and TEVcs uncaging (+heat, +light). (D) Quantification of experiments in panel D. Each dot in the chart indicates the number of Basin-4 neurons expressing mCherry in one larva. Error bars, Standard error of the mean value. Sample numbers indicated with the dots in each chart. n = 8-20. One-way ANOVA with Tukey’s two-group comparisons. ****p value<0.0001; ***p value<0.001; ns, p value> 0.05. Scale bar: 50 μm.

For the initial test in *Drosophila*, we expressed the cytoFLARE1.0 transgenes in Basin-4 neurons, which are a type of second-order neurons (SONs; i.e., postsynaptic to the nociceptors) in the nociceptive pathway ^35,36^. Heat at 39 ^°^C was applied to the third-instar larvae as a noxious cue ^37-39^, along with blue light for activating cytoFLARE (Supplementary Fig. 8). After stimulation, the larvae were raised in dark for another 16 hours to allow the transcription and translation of the mCherry reporter. In this initial test, mCherry was observed even without heat or blue light (Supplementary Fig. 8). In parallel, the control transgenic flies expressing all the cytoFLARE1.0 components except the CaM-TEVp did not express mCherry. This result suggests that the background mCherry expression from cytoFLARE1.0 was due to cleavage by co-expressed TEVp, rather than mis-trafficking of the TF component or endogenous protease cleavage. The light– and calcium-independent protease cleavage was likely caused by the overexpression of cytoFLARE1.0 transgenes throughout the larval development.

To reduce the background accumulation of the reporter protein throughout the larval development, we restricted the expression of cytoFLARE components to a narrower period by using a temperature-sensitive GAL80 transgene (GAL80^ts^) ^40,41^. The yeast protein GAL80 inhibits the GAL4-UAS system, and a temperature-sensitive mutant, called GAL80^ts^, allows temporal controls of gene expression in the GAL4-UAS system. GAL80^ts^ can inhibit GAL4 at a permissive temperature (e.g., 18 °C) but becomes inactive at a restrictive temperature (e.g., 31 °C). As a result, we can control the temperature of the environment in which the *Drosophila* live to restrict the expression timing of the UAS-transgenes. We kept transgenic larvae at 31 ^°^C for different periods ranging from 60 to 114 hours. Both 85 and 114 hour-incubation led to sufficient cytoFLARE1.0 expression and significant mCherry reporter expression upon 5-minute light and noxious heat stimulation (Supplementary Fig. 9). Since the 114-hour incubation generated higher background in the dark condition, we proceeded to characterize cytoFLARE1.0 with 85-hour GAL4-UAS expression.

Next, we tested the sensitivity of cytoFLARE1.0 for recording neuronal activity in response to noxious heat (39 °C) with various pulses (Figure 3B). Notably, we observed a significant increase in the number of Basin-4 neurons expressing mCherry by 3 and 5 pulses, but not 1 pulse, of heat stimuli (Figure 3C). This result suggests that cytoFLARE activation is neuronal activity-dependent but its sensitivity to more transient calcium response might need further improvement.

We also investigated the blue light intensity required for activating cytoFLARE1.0 in *Drosophila* larvae. As demonstrated in Supplementary Fig. 10, the blue light intensity of 100 and 200 μW/mm^2^ induced a significant increase in light– and heat-dependent signals, while 50 μW/mm^2^ did not. Therefore, we proceeded with the light intensity of 100 μW/mm^2^ for optimal TEVcs uncaging in *Drosophila*.

While noxious heat can be applied on the order of minutes, optogenetic stimulation enables us to evaluate cytoFLARE on the order of seconds. Moreover, optogenetic stimulation of upstream or presynaptic neurons is commonly used for circuit and synaptic studies. Thus, in addition to heat as a natural cue, we also investigated if cytoFLARE1.0 could be activated by optogenetic stimulation of presynaptic neurons. To this end, we expressed CsChrimson, a red-light sensitive optogenetic actuator ^42^, in larval nociceptors with a QF driver ^43-45^, based on the previously established cytoFLARE1.0 transgenic *Drosophila* (Figure 4A). *Drosophila* larvae were exposed to red light with 5 s on/ 5 s off flashes for 3 pulses (1 min flashes, 1 min off each pulse) for stimulation and synchronous blue-light flashes for TEVcs uncaging (Figure 4B). We investigated red light intensities ranging from 20 to 200 μW/mm^2^, and found that when the red light intensity was 35 μW/mm^2^ or higher, there were significant increases in the number of mCherry-positive Basin-4 neurons in larvae exposed to both red and blue light, compared to those exposed to only red light, only blue light, or neither (Figure 4C and Figure 4D). However, there is a lack of significant increase in mCherry-positive neurons under weaker optogenetic stimulations (i.e., red light intensity at < 20 μW/mm^2^). Overall, the number of mCherry-positive cells increased significantly as the red light intensity increased from 20 μW/mm^2^ to 50 μW/mm^2^ and saturated at 50 μW/mm^2^ (Figure 4E). These results demonstrate that cytoFLARE1.0 can be activated by the optogenetic stimulation in a wide dynamic range.

**Figure 4.**
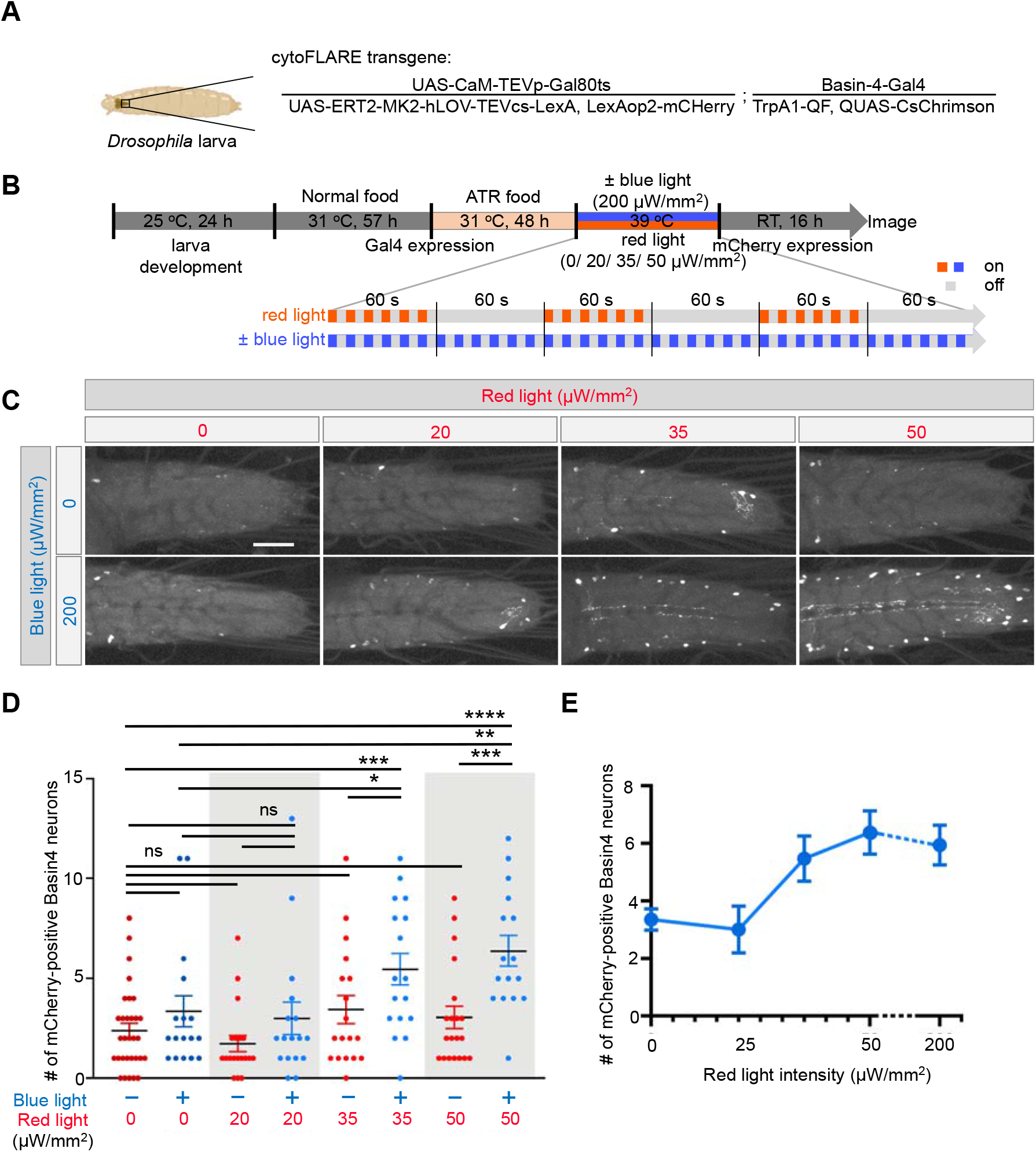
*In vivo* characterization of cytoFLARE1.0 in response to optogenetic stimulation in *Drosophila*. (A) Schematic of a larva expressing cytoFLARE1.0 in Basin-4 neurons and CsChrimson in the thermal nociceptors. (B) Experimental workflow for characterizing cytoFLARE1.0 in response to optogenetic stimulation in *Drosophila*. At early 3^rd^ instar, larvae were kept at 31°C for 105 h to inactivate GAL80^ts^ so that the TEVp and ER^T2^-transcription factor transgenes could be expressed. The larvae were transferred from normal food to ATR-containing food 48 h before stimulation to allow proper folding and membrane insertion of CsChrimson The larvae were exposed to pulsed red light (60s stimulation + 60s rest, 3 times, and the red light flashed in a cycle of 5 s on and 5 s off during stimulation) of different intensity and 200 μW/mm^2^ blue light (flashed in 5 s on/ 5 s off cycles, synchronized with red light) or no blue light. Subsequently, the larvae were kept at room temperature for 16 h to allow the expression of the mCherry fluorescent reporter. (C) Representative fluorescence images of Basin-4 neurons expressing mCherry in the condition shown by left and upper sides. (D) Quantification of experiments in panel C. Each dot in the chart indicates the number of Basin-4 neurons expressing mCherry in one larva. Error bars, Standard error of the mean value. Sample numbers indicated with the dots in each chart. n = 8-20. One-way ANOVA with Tukey’s two-group comparisons. ****p value<0.0001; ***p value<0.001; **p value<0.01; *p value<0.05; ns, > 0.05. Scale bar: 50 μm. (E) Changes in mean numbers of mCherry-positive cells in conditions of blue light plus different intensity of red light. Error bars, Standard error of the mean value.

### Apply cytoFLARE to manipulate neuronal activity, based on the activity history of the neuron

While recording neuronal activity with FLARE systems allows us to identify the neurons that are activated during a cognitive or behavioral process, the ability to manipulate the activated neurons is required to determine the essential role of these activated neurons involved in the process. In theory, the FLARE system also offers a way to manipulate neuronal activity, based on the previous activity levels of these neurons. However, this potentially important application of FLARE has not been demonstrated in *Drosophila*. To fill in this gap, we tested in the larval nociceptive system the possibility of using cytoFLARE to manipulate the activity of neurons based on their previous activity levels.

A08n neurons are one of the ideal cell types for assessing cytoFLARE1.0’s ability of optogenetic re-activation in *Drosophila*. In the larval nociceptive system, the A08n neurons (Figure 5A) are the most sensitive type of SONs to nociceptor activation ^38^. There are only one A08n neuron on the left and one on the right side of the larval ventral nerve cord (VNC) ^35^. We demonstrated that optogenetic activation of A08n neurons by expressing the actuator CsChrimson ^42^ with the GAL4/UAS system increased larvae’s avoidance response to noxious heat, including the count and duration of body curling and the probability of rolling (Figure 5A and 5B), as previously reported ^35,38,39^. The video-recordings of behavior were automatically analyzed by the artificial-intelligence-based software LabGym 2.2 ^46,47^.

**Figure 5.**
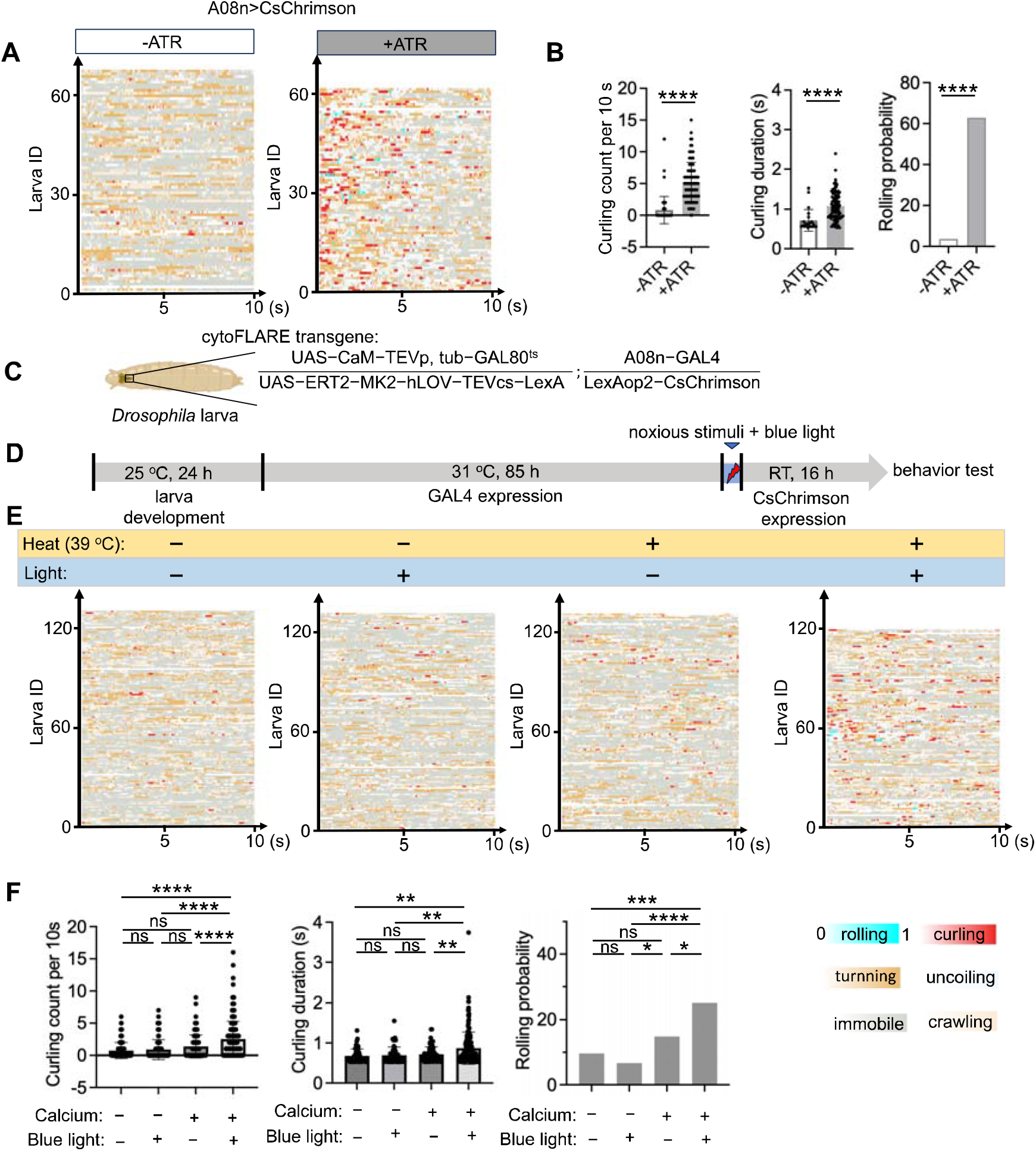
Manipulation of neuronal activity by cytoFLARE1.0 in *Drosophila*. (A) Raster plot of directly stimulate A08n neurons by optogenetics, stimulation onset from 0, stimulation duration: 10 s; Color bars: the same with panel C, stimulation onset from 0, stimulation duration: 10 s. (B) Quantification of curling count, curling duration, and rolling probabilities calculated by the LabGym 2.2 for panel C. Un-paired Student T test. Error bars, Standard error of the mean value. Sample numbers indicated with the dots in each chart. (C) Schematic of a larva expressing cytoFLARE1.0 in A08n neurons with a temperature-sensitive Gal80 transgene. (B)Experimental workflow for using cytoFLARE1.0 to manipulate A08n neuronal activity in *Drosophila*. After egg collection for 24 hours, larvae were kept at 31°C for 85 h to inactivate GAL80^ts^ so that the TEVp and ER^T2^-transcription factor transgenes could be expressed. The larvae were then exposed to 5 minutes of noxious heat stimulation (39°C) and blue light (450 nm, 100 μW/mm^2^) or one of the two treatments. Subsequently, the larvae were kept at room temperature for 16 h to allow the expression of the CsChrimson reporter. (D) Raster plots generated by LabGym 2.2 showing the frame-wise categorizations for 4 groups ((1) with neither noxious stimulation nor TEVcs uncaging (-heat,-light); (2) with noxious stimulation but not TEVcs uncaging (+heat, −light); (3) without noxious stimulation but with TEVcs uncaging (-heat, +light); and (4) with both noxious stimulation and TEVcs uncaging (+heat, +light)) after nociceptors were optogenetically activated. Stimulation onset: 0; stimulation duration: 10 s; Color bars: behavioral categories (color intensities indicate the behavioral probabilities). The x axis represents the time and y axis represents the larval ID. (E) Quantification of curling count, curling duration, and rolling probabilities calculated by the LabGym 2.2 for panel C. One-way ANOVA with Tukey’s two-group comparisons. ****p value<0.0001; ***p value<0.001; **p value<0.01; *p value<0.05; ns, p value> 0.05.

After confirming CsChrimson’s effect in A08n neurons, we tested cytoFLARE1.0 by expressing the cytoFLARE1.0 transgenes in A08n neurons, along with the CsChrimson as the reporter gene (Figure 5C). To activate cytoFLARE1.0, we stimulated larval nociceptors with noxious heat (39 °C, 5 min) and 450-nm blue light (100 μW/mm^2^) (Figure 5D). After stimulation, the larvae were raised in dark for another 16 hours to allow the transcription and translation of CsChrimson to take place. Subsequently, we stimulated CsChrimson with 617 nm red light (100 μW/mm^2^) and recorded the behavioral response. We tested 4 conditions: (1) no noxious heat or TEVcs uncaging (“-heat, –light”); (2) with noxious heat but not TEVcs uncaging (“+ heat, −light”); (3) without noxious heat but with TEVcs uncaging (“-heat, + light”); and (4) with both noxious heat and TEVcs uncaging (“+ heat, + light”). Compared with the control groups, larvae in the “+ heat, + light” group showed an increase in the curling count and duration as well as the probability of body rolling, which are characteristic nociceptive responses (Figure 5E and 5F). These results suggest that cytoFLARE1.0 not only reports the activity of neurons during sensory stimulation but also allows reactivation of those previously activated by the sensory stimulation.

## Discussion

Activity-dependent labeling and manipulation of neurons activated during specific behavior are important for dissecting the functional neuronal circuits. In this work, we present an improved FLARE reporter, cytoFLARE1.0. cytoFLARE1.0 uses ER^T2^ fusion to localize the TF component in the cytosol and exclude it from the nucleus, facilitating its interaction with the cytosol-localized CaM-TEVp component. We also incorporated a higher affinity CaM-MK2 pair in cytoFLARE1.0 to increase its calcium sensitivity and the fast-reset LOV domain for more precise temporal gating resolution. Overall, cytoFLARE1.0 shows higher activation efficiency than FliCRE.

We further established the application of cytoFLARE1.0 in *Drosophila* larvae. We found that restricted expression of cytoFLARE1.0 components to 2-3 days before the experiment, using the GAL4-GAL80 transcriptional system, is necessary to reduce cytoFLARE1.0’s background activation. We then demonstrated cytoFLARE1.0 can label Basin-4 neurons activated by 3 pulses of 1-minute noxious heat stimuli, along with blue light co-stimulation to uncage the protease cleavage site. CytoFLARE1.0 can also label Basin-4 neurons activated by optogenetic stimulation of their presynaptic neurons, the nociceptors. Further, we showed that the A08n neurons can be reactivated via cytoFLARE1.0 to drive the noxious response in the *Drosophila larvae*.

We have demonstrated for the first time of time-gated neuronal-activity-dependent labeling and manipulation of neurons in *Drosophila*. This cytoFLARE1.0-based system fills the current gap of tools for labeling activated neurons in *Drosophila* with temporal gating and will improve the dissection of neuronal circuits and their functions in *Drosophila*. Additionally, the *Drosophila* larva provides a powerful platform for testing cytoFLARE and other time-gated calcium integrators. For example, light can be directly applied to the larvae without the need of optical fiber implantation. Moreover, expression levels through transgenes are more consistent than those through mixed viral injections.

There are also many remaining challenges associated with using cytoFLARE1.0 to label activated neurons in living organisms. First, as a multi-component reporter system, cytoFLARE1.0 is highly dependent on protein expression levels; therefore, fine-tuning of the expression levels of the various components will be required for optimal performance in different neuronal circuits and living organisms. Additionally, the sensitivity of cytoFLARE1.0 is still limited as minutes of stimulation are needed in primary rat cultured neurons and *Drosophila* larvae. cytoFLARE1.1 with a faster-kinetics protease could potentially further improve the sensitivity in living organisms in the future.

In summary, as an improved calcium integrator, cytoFLARE1.0 can label active neuronal ensembles with time sensitivity of minutes and enable further investigation of the causal relationship between the specific neuron ensembles and animal behavior. Although this work focuses on *Drosophila* applications, cytoFLARE can also be potentially adapted for applications in other organisms.

## Acknowledgments

This research was supported by National Institutes of Health (NIH) to B.Y. (R01NS128500) and W.W. (R01DA053200), and National Science Foundation to W.W. (CHE-2235835). *Drosophila* Stocks from the Bloomington *Drosophila* Stock Center (NIH P40OD018537) were used in this study. The content is solely the responsibility of the authors and does not necessarily represent the official views of NIH or NSF.

## Author Contributions

G.Z. and W.W. conceived the reporter designs and experimental designs for HEK293T cell testing and for primary neuron testing. R.L. and B.Y. conceived the experimental designs for *Drosophila* testing. G.Z. and W.W.W. performed HEK293T cell experiments. G.Z. and A.B. performed primary neuron experiments. R.L., Y.M., J.M., and Y.H. performed *Drosophila* experiments. G.Z. and W.W.W. cloned the constructs and prepared viruses used in these experiments. G.Z., R.L., Y.M., B.Y., and W.W. analyzed data, and wrote and edited the manuscript.

## Declaration of Interests

The authors declare no competing interests.

## Supplemental information

Document S1. Figures S1–S10.

## Materials and Methods

### Cloning

Constructs for protein expression in HEK293T cells were cloned into the ampicillin-resistant pLX208 lentiviral vector for infection. Constructs for neuron experiments were cloned into the ampicillin-resistant pAAV viral vector with the synapsin promoter. Constructs for making transgenic *Drosophila* larvae were cloned into the ampicillin-resistant pUASTattB vector.

For PCR fragment amplification, Q5 polymerase was used to amplify fragment. The vectors were double-digested with NEB restriction enzymes and ligated to gel-purified PCR products by Gibson assembly. Heat shock transformation was performed to introduce the ligated plasmid products into XL-1-blue bacteria.

### HEK293T cell culture, lentivirus production and infection

HEK293T cells with a passage number below 20 were cultured at 37 °C under 5% CO_2_ in complete growth media composed of 1:1 DMEM (Dublecco’s Modified Eagle Medium, Gibco): MEM (Modified Eagle Medium, Gibco), 10% FBS (Fetal Bovine Serum, Sigma), and 1% Pen Strep (Gibco).

For lentivirus production of each construct, a T25 flask was pre-treated with 1 mL of 20 μg/mL HFN at 37 °C for at least 10 min. After aspiration of HFN, HEK293T cells were plated at 70-90% confluence. 2.5 μg viral DNA, 0.25 μg pVSVG, and 2.25 μg delta8.9 lentiviral helper plasmid were combined with 250 μL of DMEM and thoroughly mixed. 25 μL PEI MAX was then added to the mixture. After 15 min incubation at room temperature, 1 mL complete growth media was added to the DNA-PEI mixture and transferred to the T25 flask. After incubation at 37 °C under 5% CO_2_ for 36-48 h, the lentivirus supernatant was collected, flash frozen in liquid nitrogen, and stored at –80 °C for up to one year.

For lentivirus infection in each well of the 48-well plastic plates, a mixture of 150 μL TEVp, transcription factor, and UAS-reporter gene lentiviruses (50 uL respectively) was prepared and then well mixed with 200 μL of confluent HEK293T cells by pipetting. The cell lentivirus mixture was transferred to each well of the 48-well plate and the cells were incubated for 36-48 h before stimulation and imaging. For lentivirus infection in 96-well white plates, all reagents were scaled 2-fold down. For lentivirus infection in 24-well plastic plates, all reagents were scaled 2-fold up.

### HEK293T cell stimulation for light– and calcium-dependent protease cleavage

After lentivirus infection, HEK293T cells were stimulated by 5 mM CaCl_2_ (DOT Scientific, Cat no. DSC20010) in the presence of 2 μM ionomycin (MilliporeSigma, Cat no. 407953) with blue light (470 nm) for 1 or 5 min. Cells were then washed twice with the complete growth media and incubated at 37 °C for 24 h before live-cell fluorescence imaging, luminescence reading, or flow cytometry analysis.

### Flow cytometry characterization of cytoFLARE activation efficiency

For flow cytometry analysis, HEK293T cells were plated in 24-well plastic plates. After stimulation and overnight incubation, the media was aspirated and replaced with 80 μL trypsin. The trypsinized cells were then mixed with 500 μL complete growth media and transferred to a 5 mL polystyrene round-bottom tube. LSR Fortessa cell analyzer flow cytometer (BD Biosciences) equipped with 488 nm laser and 530/30 emission filter as well as 561 nm laser and 586/15 emission filter and FlowJo was used for FACS analysis.

### HEK293T luminescence assay and data analysis

For luminescence assays, HEK293T cells were plated in 96-well white plates. After stimulation, cells in each well were incubated in 66uL complete growth media at 37 °C for 24 h. Before luminescence measurement, the plate was cooled down to the room temperature for 5 min. 66 uL Bright-Glo (Promega, Cat no. PRE2620) was then added to each well. Luminescence was immediately measured on a BioTek Cytation 5 cell imaging multimode reader using a 1 s acquisition time and gain at 150.

Three technical replicates (wells) were performed per condition and the luminescence intensity of each conditioned well was subtracted by the background determined by the well without cells. Each well was processed and plotted as a single dot. The Prism Graph Pad software was used to calculate the mean and standard error of the mean (SEM) for each condition and the unpaired two-tailed Student’s t-tests for significance.

### Rat neuronal cultures

E18 Sprague Dawley Rat Whole Brain (TransnetYX, Cat no. SDEWB) was dissociated with the Cys-activated papain solution, consisting of 2 mL L-Cysteine (Sigma, Cat no. 30089) and 100 μL papain (Sigma, Cat no. P3125), at pH 6.5. Following dissociation, 2×10^5^ neuronal cells were plated in each well of the 24-well plate pre-coated with 400 uL of poly-D-lysine (0.1 mg/mL, Sigma, Cat no. P7280) and laminin (3 μg/mL, Sigma, Cat no. L2020) premixed solution. Neuronal cells were cultured at 37 °C under 5% CO_2_ in plating medium composed of 3:1 complete neurobasal media (NGM): glial enriching medium (GEM) mixture. The NGM consists of Neurobasal Plus Medium (Gibco), 2% B27 supplement (Thermo Fisher Scientific), 1% GlutaMAX (Gibco), and 1% Pen Strep (Gibco). The GEM consists of HEK293T complete growth media, 2% B27 supplement, and 1% GlutaMAX. To maintain the neuronal culture, 300 μL old media was replaced with 500 μL fresh NGM every 4 days.

### Viral transduction of rat neuronal cultures

At DIV6, virus mixtures were prepared by mixing 210 μL of TEVp, transcription factor, and UAS-reporter gene viruses (70 μL respectively) for neurons in each well of the 24-well plate. Following buffer exchange with fresh NGM through an Amicron centrifugal filter (100KDa, MilliporeSigma), 500 μL NGM was mixed with 10-20 μL filtered virus mixture and added to neurons in each well. After viral transduction, the plate was wrapped with aluminum foil and incubated at 37 °C under 5% CO_2_.

### Neuronal culture stimulation for light– and calcium-dependent protease cleavage

At DIV12 (6 days post transduction), neurons were stimulated by 20 μM bicuculline (Sigma, Cat no. 14340) or electrical stimulation with blue light (470nm) for 1 and 5 min. For neurons treated with bicuculline, cells were washed twice with NGM and incubated at 37 °C for 24 h before live-cell fluorescence imaging.

### HEK293T and neuronal culture fluorescence imaging and data analysis

HEK293T cells and neuronal cells were imaged by a Nikon inverted confocal microscope with the 20x air objective. The microscope is outfitted with a Yokogawa CSU-X1 5000RPM spinning disk confocal head, Ti2-ND-P perfect focus system 4, and a compact 4-line laser source with the following lasers: 405 nm (100 mW), 488 nm (100 mW), 561 nm (100 mW), and 640 nm (75 mW). The combinations of laser excitation and emission filters were used as follows: EGFP (488 nm excitation; 525/36 emission); mCherry (568 nm excitation; 605/52 emission). All images were taken with a 1 s acquisition time, 50 % laser intensity, and an ORCA-Flash 4.0 LT+sCMOS camera. All images were collected and processed using the Nikon NIS-Elements software.

For HEK293T cell experiments, 3 technical replicates (wells) were performed per condition and 8 fields of view were taken per well. NIS-Elements General Analysis 3 software was used for fluorescence intensity analysis. To obtain the mean intensity for each field of view, a threshold was set to be above the autofluorescence. The mean intensity value was then multiplied by the object areas to obtain a sum intensity value. The mean of sum fluorescence intensity in the 8 fields of view from the same well was calculated and plotted as a single dot. Prism Graph Pad software was used to calculate the mean and standard error of the mean (SEM) for each condition and the unpaired two-tailed Student’s t-tests for significance.

For neuronal culture experiments, 3 technical replicates were performed per condition and 12 fields of view were taken per well. The sum intensities of mCherry and EGFP for each entire field of view were measured on NIS-Elements General Analysis 3 software and then subtracted by the autofluorescence of the plates. The mean of sum fluorescence intensity in the 12 fields of view from the same well was calculated and plotted as a single dot. Prism Graph Pad software was used to calculate the mean and standard error of the mean (SEM) for each condition and the unpaired two-tailed Student’s t-tests for significance.

### *In vivo* characterization of cytoFLARE in *Drosophila*

Both male and female foraging third-instar larvae were used. To test cytoFLARE for recording neuronal activity in *Drosophila*, cytoFLARE transgenes (UAS−CaM−TEVp, UAS−ERT2−MK2−hLOV−TEVcs−LexA) were expressed in Basin-4 neurons with the GMR57F07-GAL4. The temperature-sensitive GAL80^ts^ transgene was expressed via a tubulin promoter. The LexAop2-mCherry transgene served as the reporter for the presence of nuclear LexA. After the embryos developed into larvae, they were kept at 31°C for 60-114 h to express the cytoFLARE transgenes. Larval nociceptors were intensely stimulated with heat (39 °C). A heat-plate system was employed for heat stimulation of larvae ^39,48^. This system comprises a device capable of rapid increase in temperature from room temperature to 70°C (CP-031, TE technology, USA), and is coupled with a temperature controller (TC-720, TE technology, USA). To restrict the reporting of neuronal activity to the time-window of nociceptor stimulation, the larvae were illuminated with 450 nm blue light (50-200 μW/mm^2^). Subsequently, larvae were dissected to expose the central nervous system (CNS), followed by fixation in 4% formaldehyde for 20 minutes and imaging with a Leica SP5 confocal system.

For optogenetic stimulation of larval nociceptors, CsChrimson was expressed in the nociceptors with the TrpA1-QF transgene. After the embryos developed into larvae at 25 °C, they were kept at 31 °C for 105 h. During this period, the larvae were transferred to standard cornmeal food containing 1 mM all-trans-retinal (ATR) (A.G. Scientific) 48 h before stimulation. The larvae were then exposed to 617 nm red light (0, 20, 35, 50, 200 μW/mm^2^) to stimulate nociceptors. The red light was applied in 5s-on and 5s-off flashes for 3 pulses (1 min flashes, 1 min off each pulse). The 450 nm blue light was in 200 μW/mm^2^ and was applied in 5s-on and 5s-off flashes synchronized with red light throughout the stimulation process. Subsequently, the larvae were kept at 20-21°C for 16 h to allow the expression of the mCherry fluorescent reporter.

### Application of cytoFLARE to manipulate neuronal activity

We expressed in A08n neurons the cytoFLARE1.0 transgenes (UAS−CaM−TEVp, UAS−ERT2−MK2−hLOV−TEVcs−LexA) with the GMR82E12-GAL4. To express the optogenetic actuator CsChrimson based on A08n neuronal activity, the LexAop2-CsChrimson transgene was include in the flies. If cytoFLARE detects an increase in neuronal activity (indicated by increased calcium levels) during a specific time window (defined by blue-light illumination), the LexA transcription factor would translocate into the cell nucleus and activate the transcription of LexAop2-CsChrimson, leading to the expression of CsChrimson in A08n neurons. CsChrimson allows optogenetics activation of the A08n neurons by red-light illumination.

Embryos were collected on standard cornmeal food containing 1 mM ATR. After the egg collection for 24 h, larvae were kept at 31 °C for 85 h. Larva nociceptors were stimulated with heat (39°C, 5 min). To active the LOV domain and uncage the TEVcs, we illuminated the larvae with 450-nm blue light (100 μW/mm^2^). After stimulation, the larvae were raised in dark for another 16 hours to allow the CsChrimson gene expression. To stimulate A08n*>*CsChrimson larvae, 100 μW/mm^2^ red light (617 nm) was applied for 10 s. All LEDs used were from LUXEON StarLED. Light intensity was measured with a power meter (PM16-122, Thorlabs, USA). Larva’s response were recorded by a webcam (C910, Logitech, USA), and analyzed with the LabGym 2.2 software ^46,47^.

## Notes

### Competing Interest Statement

The authors have declared no competing interest.

