## Supplementary information for "An improved FLARE system for recording and manipulating neuronal activity"

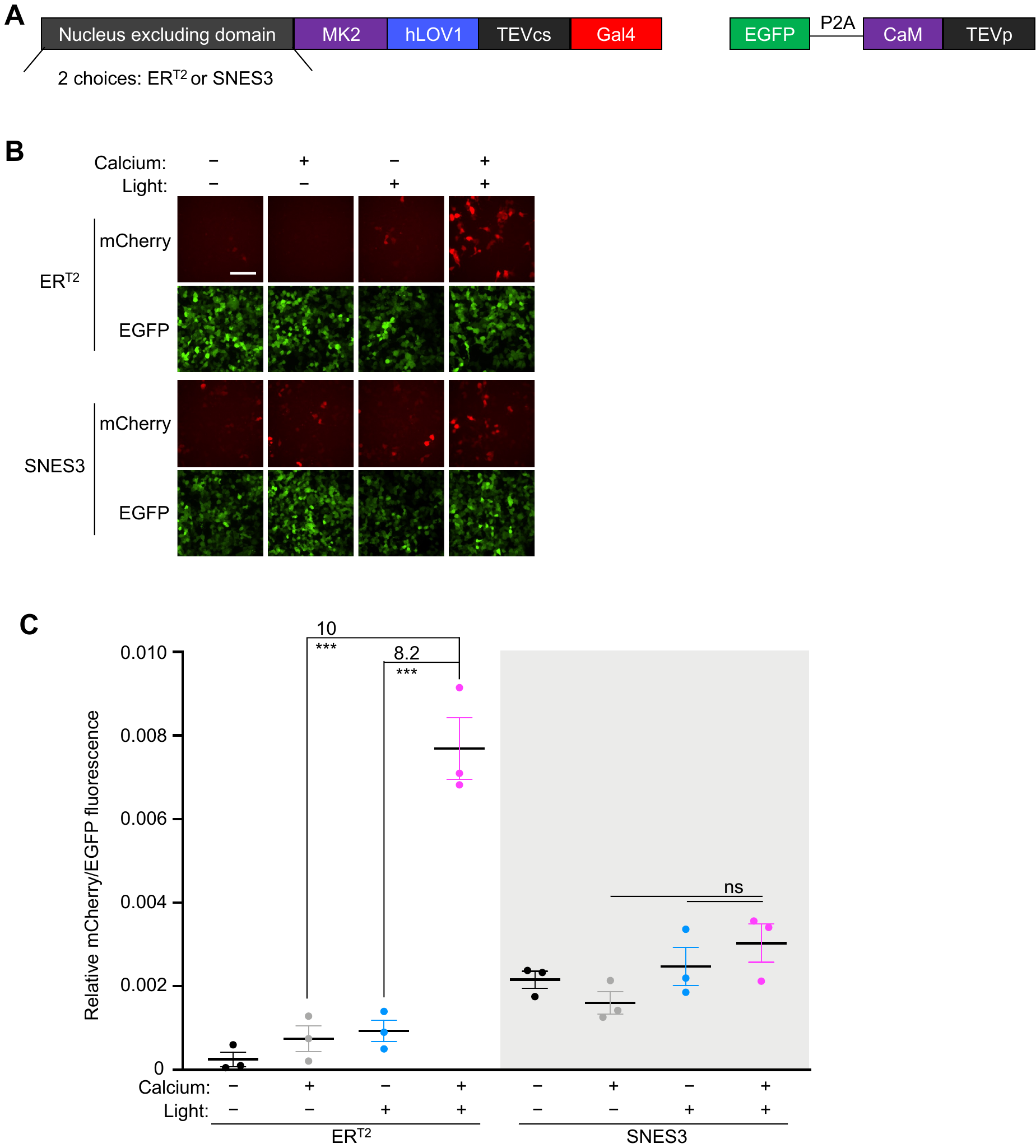

**Supplementary Figure 1. Comparison of nucleus excluding domains in cytoFLARE design in HEK293T cells.**

(A) Construct design of cytoFLARE with ER^T2^ and SNES3, respectively.

(B) Representative fluorescence images for comparing ER^T2^ and SNES3’s effect on cytoFLARE’s performance in HEK293T cells. Calcium and blue light were applied for 1 minute. mCherry, reporter gene expression of cytoFLARE. EGFP, expression marker of the cytoFLARE TEVp construct. Scale bar, 100 µm.

(C) Quantification of experiments in panel B. Values above the dots represent SBRs which were calculated by dividing the means of the conditions indicated. Error bars, standard error of the mean. The mean is represented by the thicker horizontal bar. Stars represent significance after an unpaired two-tailed Student’s t-test. n = 3. ***p value <0.001; ns, p value >0.05.

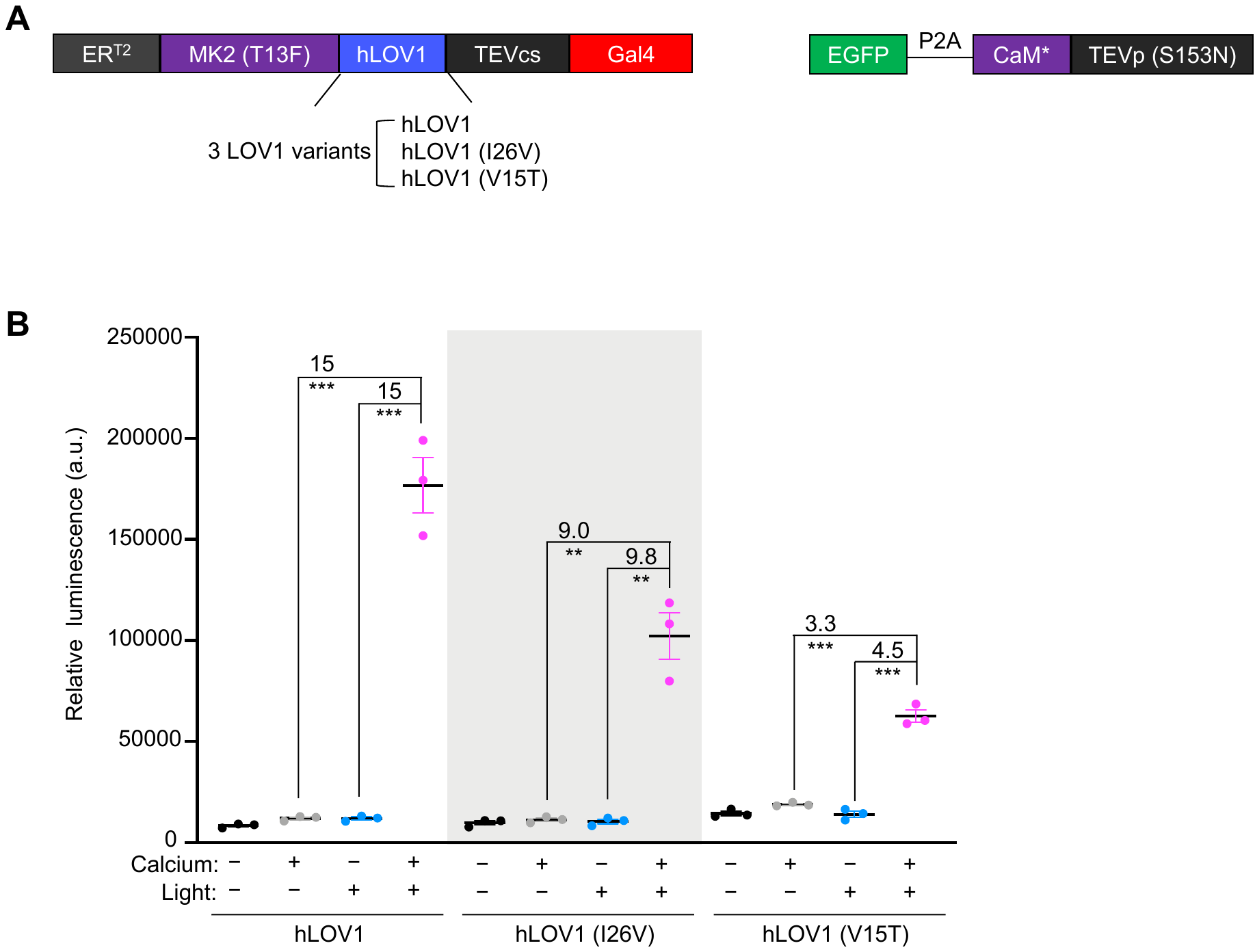

**Supplementary Figure 2. Comparison of LOV domains in cytoFLARE design in HEK293T cells.**

(A) Construct design of cytoFLARE with three different hLOV1 domains, respectively.

(B) Quantification of luminescence readout from cytoFLARE with different hLOV domains in HEK293T cells. Luciferase was used as the reporter gene. Calcium and light were applied for 1 minute. Values above the dots represent SBRs which were calculated by dividing the means of the conditions indicated. Error bars, standard error of the mean. The mean is represented by the thicker horizontal bar. Stars represent significance after an unpaired two-tailed Student’s t-test. n = 3. **p value <0.01.

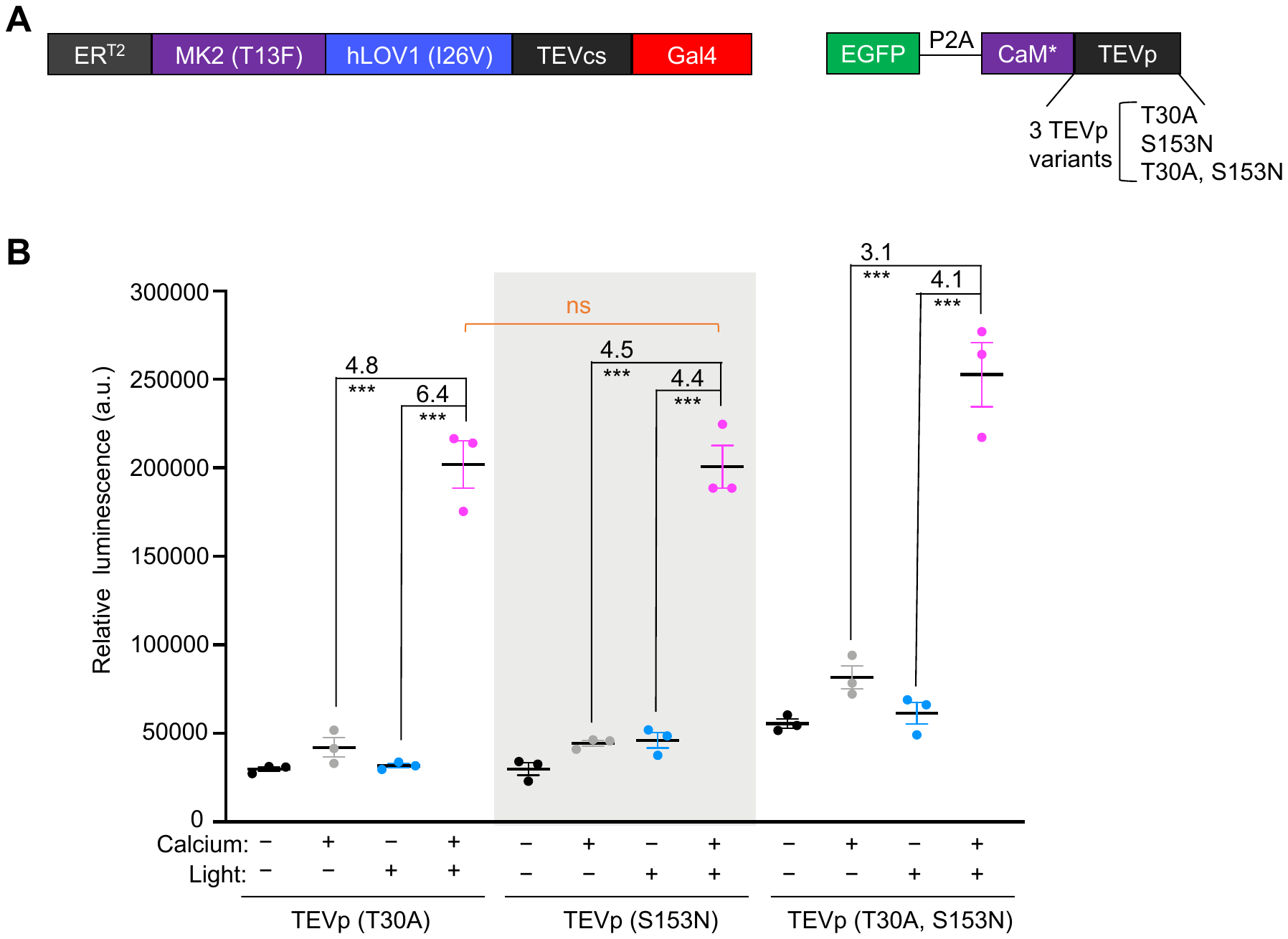

**Supplementary Figure 3. Comparison of TEVp in cytoFLARE design in HEK293T cells.**

(A) Construct design of cytoFLARE with three different TEVp, respectively.

(B) Quantification of luminescence readout from cytoFLARE with different TEVp in HEK293T cells. Luciferase was used as the reporter gene. Calcium and light were applied for 1 minute. Values above the dots represent SBRs which were calculated by dividing the means of the conditions indicated. Error bars, standard error of the mean. The mean is represented by the thicker horizontal bar. Stars represent significance after an unpaired two-tailed Student’s t-test. n = 3. ***p value <0.001; ns, p value >0.05.

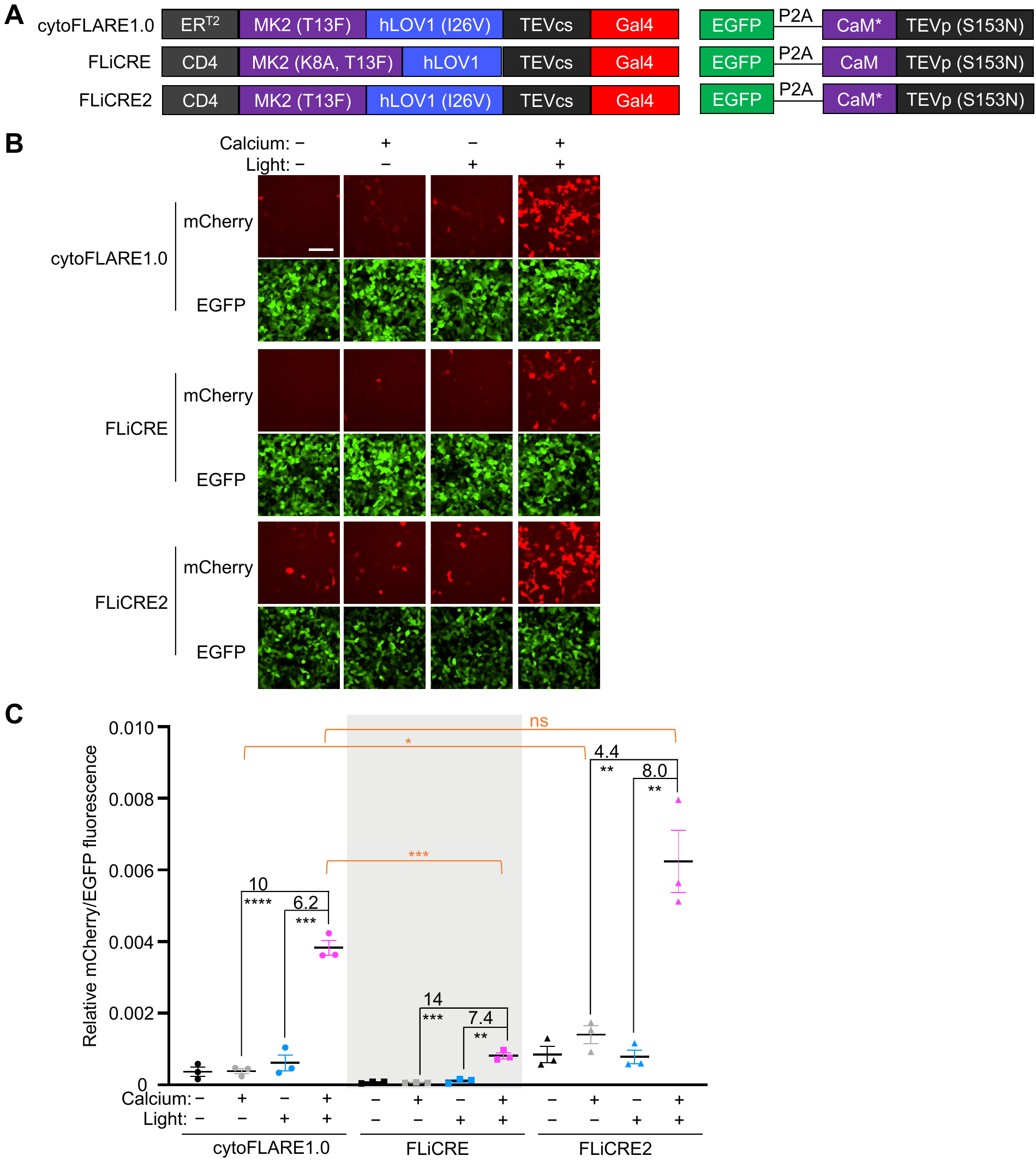

**Supplementary Figure 4. Comparison of cytoFLARE1.0 with FLiCRE and FLiCRE2 in HEK293T cells**.

(A) Construct design of cytoFLARE1.0, in comparison to FLiCRE and FLiCRE2.

(B) Representative fluorescence images for testing cytoFLARE1.0, FLiCRE and FLiCRE2 in HEK293T cells. Calcium and light were applied for 1 minute. mCherry, reporter gene expression. EGFP, expression marker of the TEVp construct. Scale bar, 100 µm.

(C) Quantification of experiments in panel B. Values above the dots represent SBRs which were calculated by dividing the means of the conditions indicated. Error bars, standard error of the mean. The mean is represented by the thicker horizontal bar. Stars represent significance after an unpaired two-tailed Student’s t-test. n = 3. ****p value <0.0001; ***p value <0.001; **p value <0.01; *p value<0.05; ns, p value >0.05.

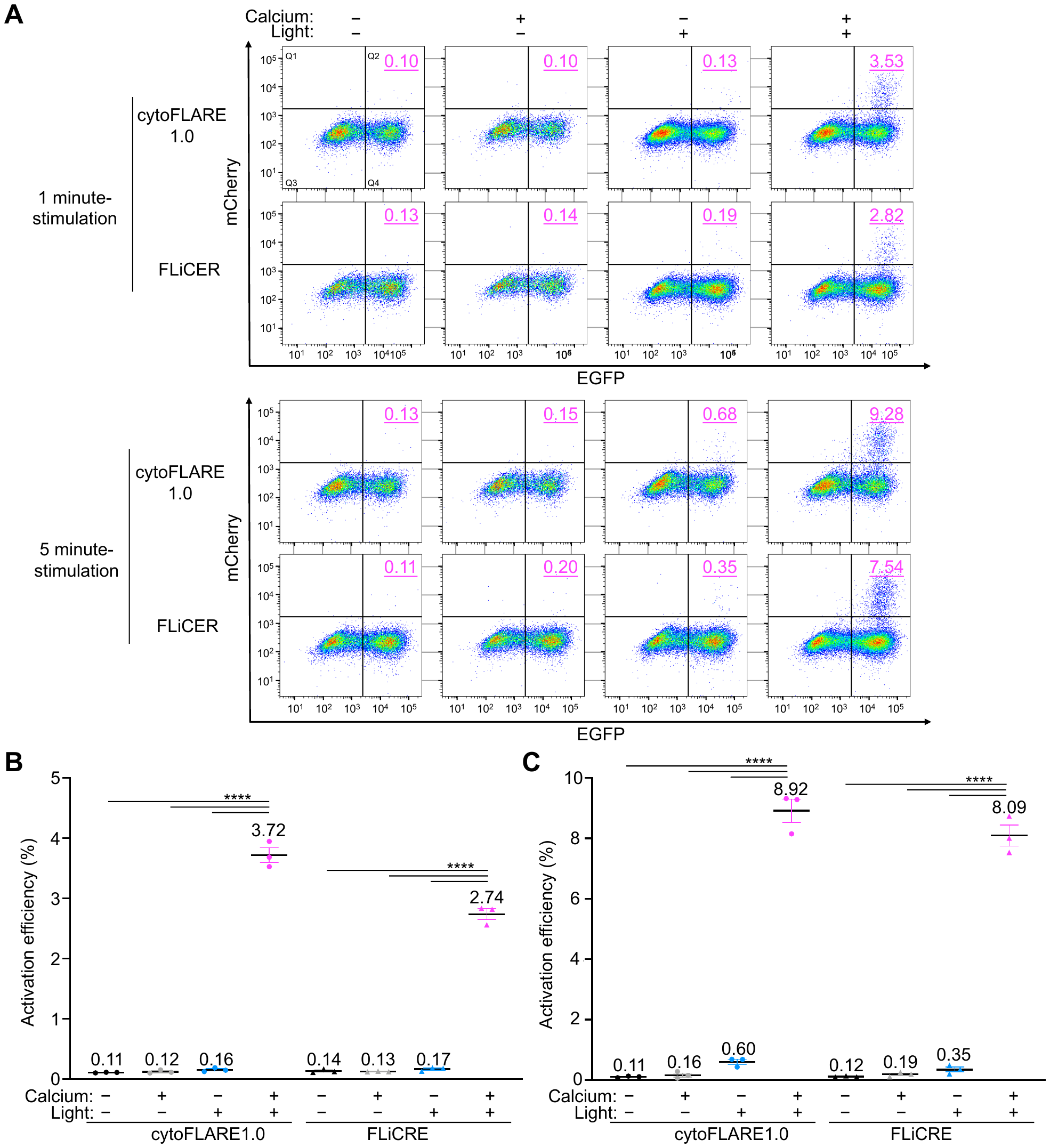

**Supplementary Figure 5. Characterization of the activation efficiency of cytoFLARE1.0 in HEK293T cells, compared to FLiCRE.**

(A) Flow cytometry analysis of HEK293T cells expressing cytoFLARE1.0 and FLiCRE, following the live cell imaging shown in Figure 1D and Figure 1E. The activation efficiency of cytoFLARE1.0 or FLiCRE was calculated by dividing the number of cells expressing the reporter gene mCherry (in Q2) divided by the total number of cells expressing the reporter (in both Q2 and Q4). The activation efficiency in each condition from 1 replicate was shown and represented by the magenta number in the top right of each FACS plot.

(B) and (C) Quantification of experiments in panel A for 1-minute stimulation and 5-minute stimulation, respectively. Values above the dots represent the mean activation efficiency in each condition from 3 replicates. Error bars, standard error of the mean. The mean is represented by the thicker horizontal bar. Stars represent significance after an unpaired two-tailed Student’s t-test. n = 3. ****p value <0.0001.

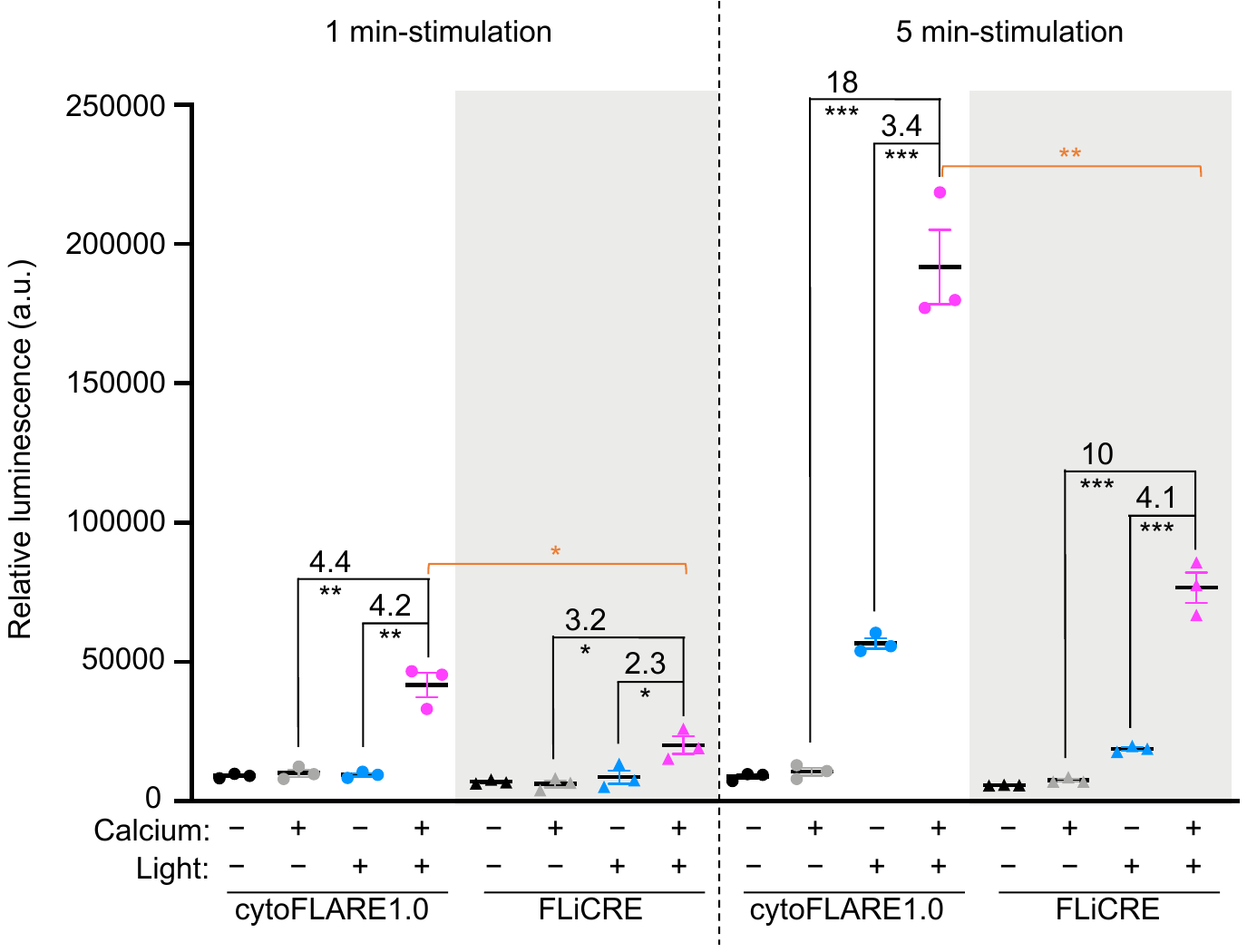

**Supplementary Figure 6. Comparison of cytoFLARE1.0 and FLiCRE with a luciferase reporter gene in HEK293T cells.** Quantification of luminescence readout from cytoFLARE and FLiCRE. Luciferase was used as the reporter gene. Calcium and light were applied for 1 minute or 5 minutes. Values above the dots represent SBRs which were calculated by dividing the means of the conditions indicated. Error bars, standard error of the mean. The mean is represented by the thicker horizontal bar. Stars represent significance after an unpaired two-tailed Student’s t-test. n = 3.  ****p value <0.0001; ***p value <0.001; **p value <0.01; *p value <0.05

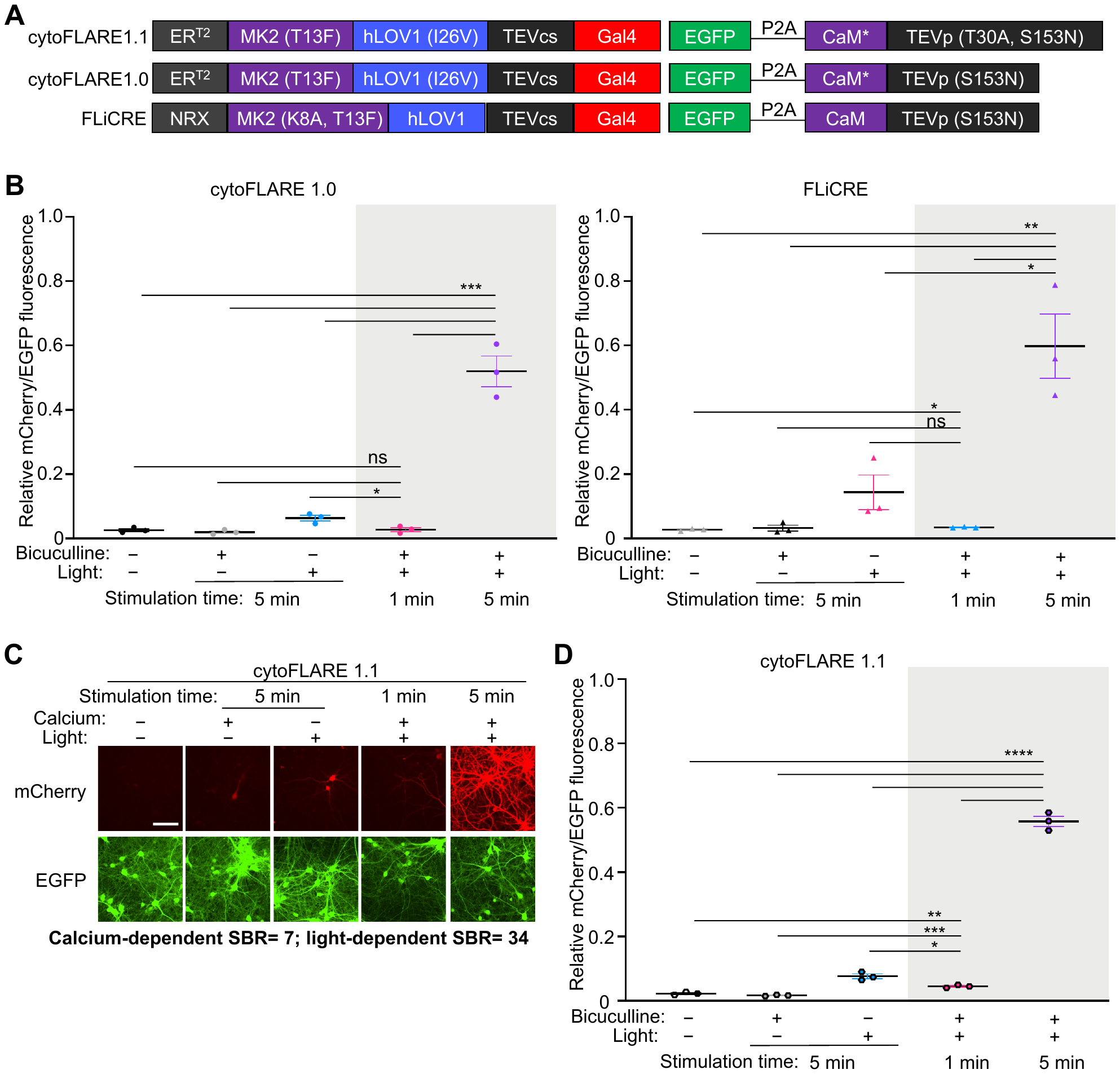

**Supplementary Figure 7. Time sensitivity characterization of cytoFLARE1.0, cytoFLARE1.1, and FLiCRE for recording neuronal activity in rat primary neurons.**

(A) Construct design of cytoFLARE1.1, in comparison to FLiCRE and cytoFLARE1.0.

(B) Quantification of fluorescence images shown in Figure 2C.

(C) Representative fluorescence images for characterizing cytoFLARE1.1 in primary rat neurons. Neurons were treated by bicuculline continuously for 1 and 5 minutes. Meanwhile, blue light was applied in pulses at 2 s on/ 4 s off for a total time of 1 and 5 minutes. mCherry, reporter gene expression. EGFP, expression marker of the TEVp construct. Scale bar, 100 µm.

(D) Quantification of experiments in panel C. Values above the dots represent SBRs which were calculated by dividing the means of the conditions indicated. Error bars, standard error of the mean. The mean is represented by the thicker horizontal bar. Stars represent significance after an unpaired two-tailed Student’s t-test. n = 3. ****p value <0.0001; ***p value <0.001; **p value <0.01; *p value<0.05; ns, p value >0.05.

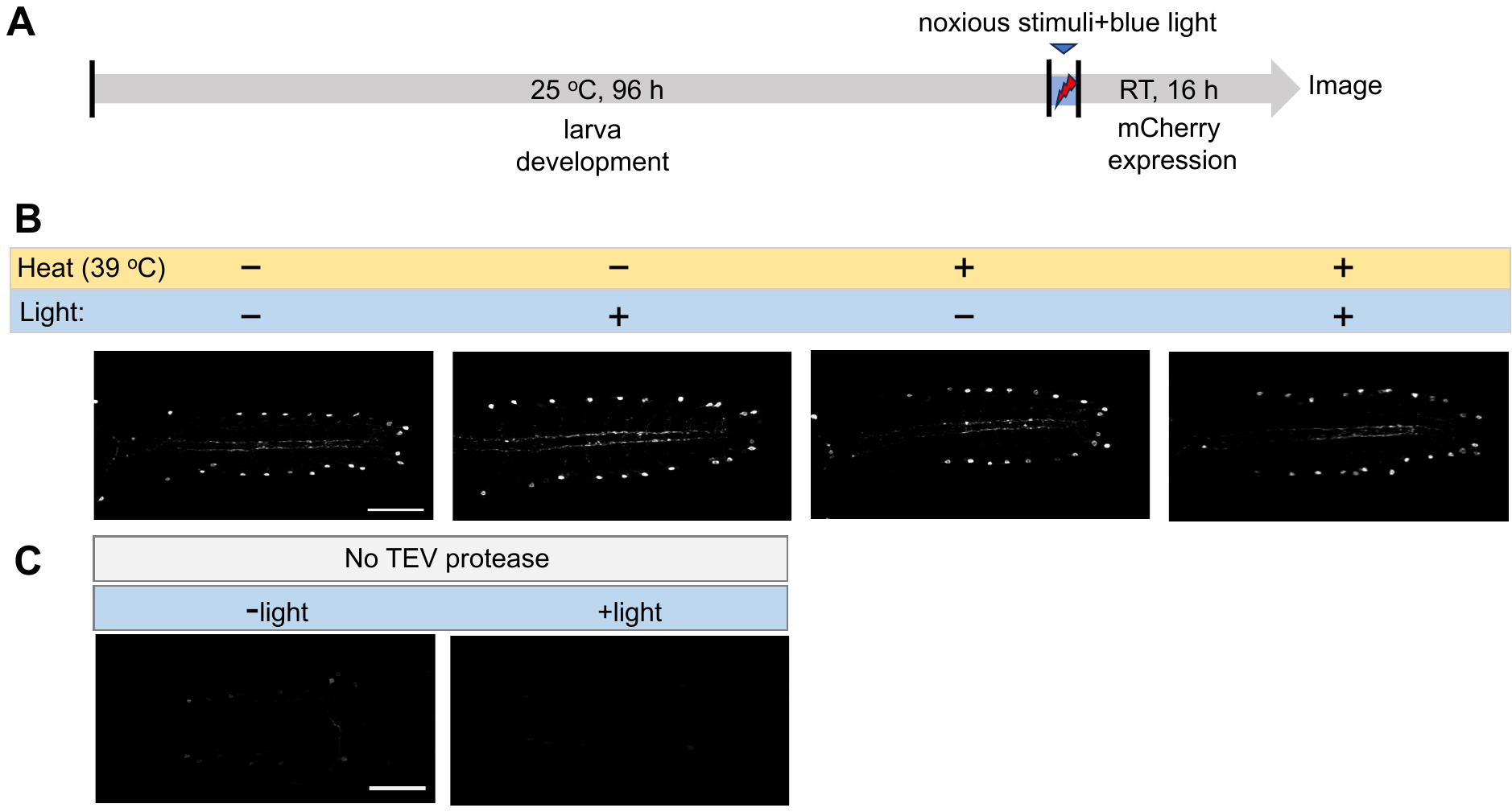

**Supplementary Figure 8. Leaky test of cytoFLARE1.0 in *Drosophila*.**

(A) Experimental workflow for testing cytoFLARE in response to noxious stimuli in *Drosophila*. At 3^rd^ instar, larvae were exposed to noxious heat stimulation (39°C, 5 min) and blue light (450 nm, 100 μW/mm^2^) or one of the two treatments. Subsequently, the larvae were kept at room temperature for 16 h to allow the expression of the mCherry fluorescent reporter.

(B) Representative fluorescence images of Basin-4 neurons expressing mCherry, which indicates the cytoFLARE activation, in the condition shown in panel. There are 4 groups: (1) with neither noxious stimulation nor TEVcs uncaging (-heat, -light); (2) with noxious stimulation but not TEVcs uncaging (+heat, -light); (3) without noxious stimulation but with TEVcs uncaging (-heat, +light); and (4) with both noxious stimulation and TEVcs uncaging (+heat, +light).

(C) Representative fluorescence of images of Basin-4 neurons expressing mCherry, which indicates cytoFLARE background in the absence of TEVp transgene. Scale bar: 50 µm.

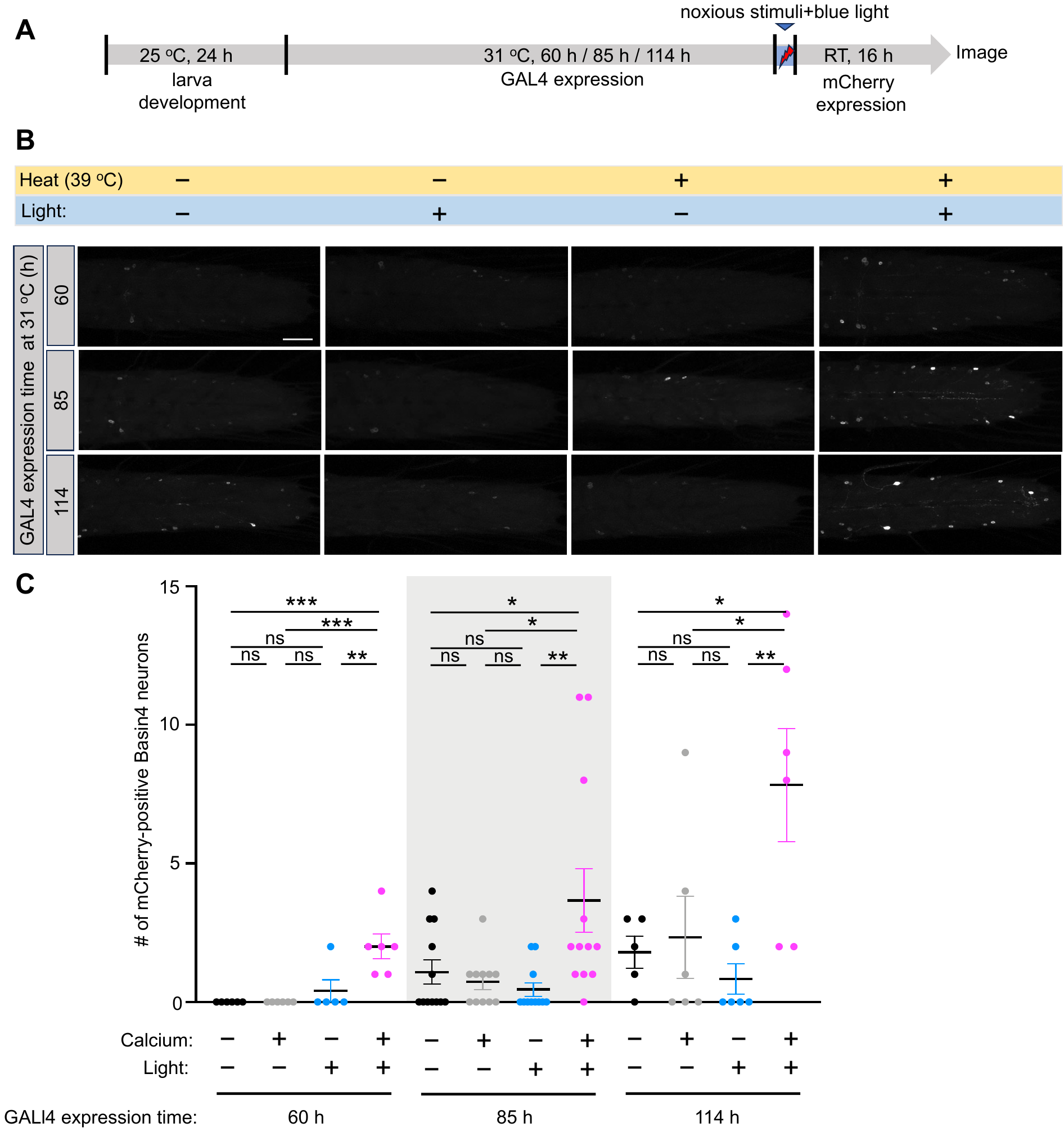

**Supplementary Figure 9. Optimization of expression of cytoFLARE1.0 in *Drosophila*.**

(A) Experimental workflow for optimizing cytoFLARE expression. At early 3^rd^ instar, larvae were kept at 31°C for 60 h, 85 h, or 114 h to inactivate GAL80^ts^ so that the TEVp and ERT2-transcription factor transgenes could be expressed. The larvae were then exposed to noxious heat stimulation (39°C, 5 min) and blue light (450 nm, 100 μW/mm^2^) or one of the two treatments. Subsequently, the larvae were kept at room temperature for 16 h to allow the expression of the mCherry fluorescent reporter.

(B) Representative fluorescence images of Basin-4 neurons expressing mCherry, which indicates the cytoFLARE activation, in the condition shown in panel A. There are 4 groups: (1) with neither noxious stimulation nor TEVcs uncaging (-heat, -light); (2) with noxious stimulation but not TEVcs uncaging (+heat, -light); (3) without noxious stimulation but with TEVcs uncaging (-heat, +light); and (4) with both noxious stimulation and TEVcs uncaging (+heat, +light).

(C) Quantification of experiments in panel B. Each dot in the chart indicates the number of Basin-4 neurons expressing mCherry in one larva. Error bars, Standard error of the mean value. Sample numbers indicated with the dots in each chart. n = 5-12. One-way ANOVA with Tukey’s two-group comparisons. ***p value<0.001; **p value < 0.01; *p value < 0.05; ns, p value > 0.05. Scale bar: 50 µm.

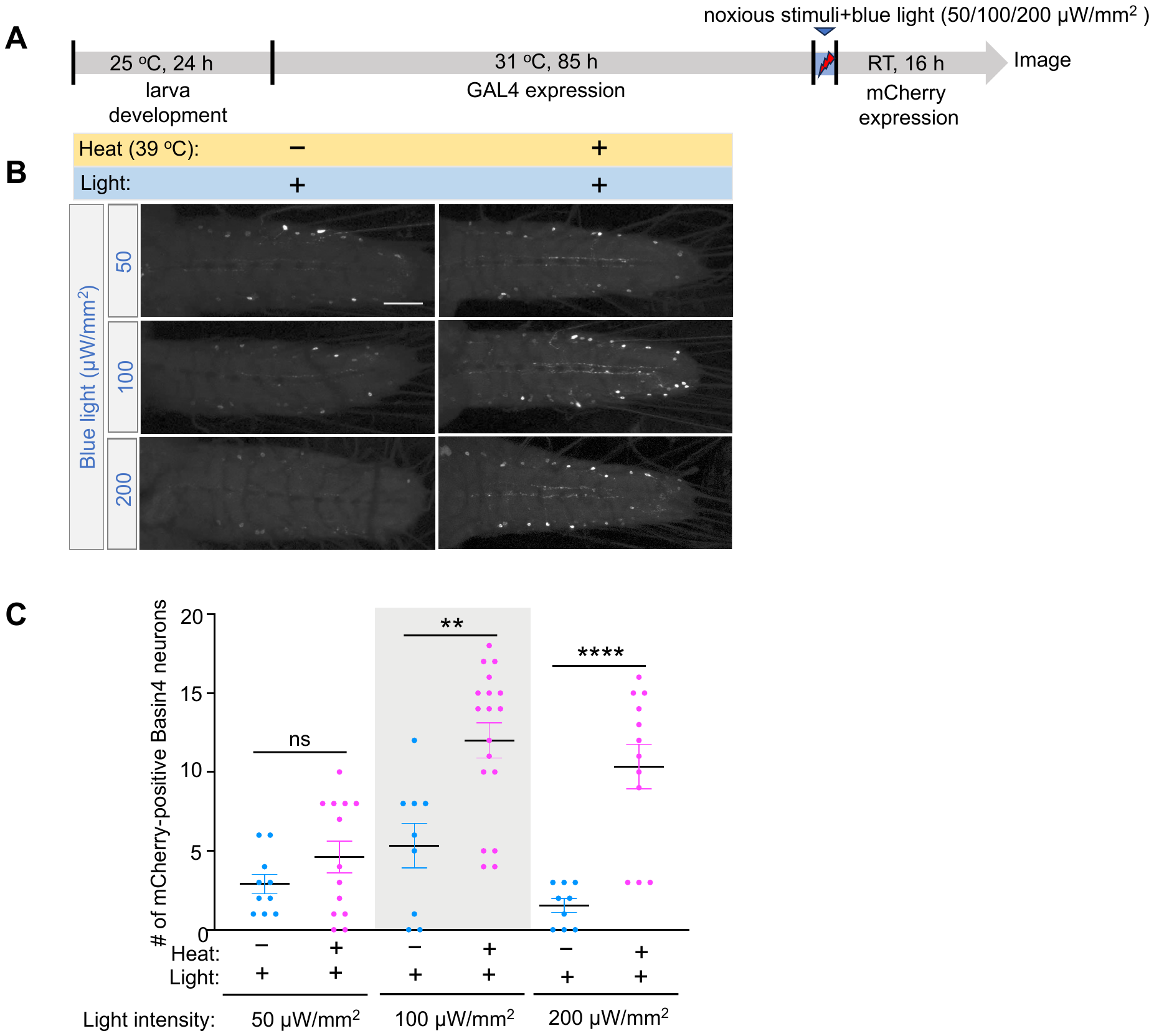

**Supplementary Figure 10. Optimization of blue light intensity for stimulating cytoFLARE.**

(A) Experimental workflow for testing cytoFLARE in response to different light intensity in *Drosophila*. At early 3^rd^ instar, larvae were kept at 31°C for 85 h to inactivate GAL80^ts^ so that the TEVp and ERT2-transcription factor transgenes could be expressed. The larvae were then exposed to noxious heat stimulation (39°C, 1 min on, 3 min off, 3 times) and blue light with different intensity (50 μW/mm^2,^ ,100 μW/mm^2^, 200 μW/mm^2^) or one of the two treatments. Subsequently, the larvae were kept at room temperature for 16 h to allow the expression of the mCherry fluorescent reporter.

(B) Representative fluorescence images of Basin-4 neurons expressing mCherry, which indicates the cytoFLARE activation, in the condition shown in panel A. There are 2 groups: (1) without noxious stimulation but with TEVcs uncaging (-heat, +light); and (2) with both noxious stimulation and TEVcs uncaging (+heat, +light).

(C) Quantification of experiments in panel B. Each dot in the chart indicates the number of Basin-4 neurons expressing mCherry in one larva. Error bars, Standard error of the mean value. Sample numbers indicated with the dots in each chart. n = 9-18. One-way ANOVA with Tukey’s two-group comparisons. ****p value<0.0001; **p value < 0.01; ns, p value > 0.05. Scale bar: 50 µm.
